# Replication dynamics of recombination-dependent replication forks

**DOI:** 10.1101/2020.07.03.186676

**Authors:** Karel Naiman, Eduard Campillo-Funollet, Adam T. Watson, Alice Budden, Izumi Miyabe, Antony M. Carr

## Abstract

DNA replication fidelity is essential for maintaining genetic stability. Forks arrested at replication fork barriers can be stabilised by the intra-S phase checkpoint, subsequently being rescued by a converging fork, or resuming when the barrier is removed. However, some arrested forks cannot be stabilised and fork convergence cannot rescue in all situations. Thus, cells have developed homologous recombination-dependent mechanisms to restart persistently inactive forks. To understand HR-restart we use polymerase usage sequencing to visualize *in vivo* replication dynamics at an *S. pombe* replication barrier, *RTS1*, and model replication by Monte Carlo simulation. We show that HR-restarted forks synthesise both strands with Pol δ for up to 30 kb without maturing to a δ/ε configuration and that Pol α is not used significantly on either strand, suggesting the lagging strand template remains as a gap that is filled in by Pol δ later. We further demonstrate that HR-restarted forks progress uninterrupted through a fork barrier that arrests canonical forks. Finally, by manipulating lagging strand resection during HR-restart by deleting *pku70*, we show that the leading strand initiates replication at the same position, signifying the stability of the 3’ single strand in the context of increased resection.

## Introduction

Each human cell has to correctly copy ∼6.5 billion bps when it divides. DNA replication is impressively accurate such that each cell division produces only a handful of errors^1^. However, numerous internal and external factors can impact on replication fidelity and cancer development is associated with increased intrinsic replication stress^2^. A number of other diseases including neurological, neurodegenerative and neuromuscular disorders (e.g. Friedreich’s ataxia, myotonic dystrophy, spinocerebrallar ataxias, fragile X syndrome and Huntington’s disease) are caused by replication-dependent slippage triggering nucleotide repeats instability^3^.

Replication forks (RFs) are disrupted by different obstacles^4^, including DNA-bound proteins, DNA damage, structure forming sequences (i.e. G-quadruplexes, inverted repeats, trinucleotide repeats) and DNA/RNA hybrids. An RF encountering such a barrier might be resolved by one of the following three strategies: (i) activation of the intra-S-phase checkpoint to stabilise the arrested RF. This preserves the replisome and the fork structure to protect the DNA from inappropriate modification. A stabilised RF can resume replication when the blockage is resolved, for example by the activity of DNA repair or accessory helicases. Alternatively, a stabilised fork can be resolved by an incoming replication fork (termination). However, not all arrested forks expose sufficient single-stranded DNA to activate the intra-S phase checkpoint and such are said to “collapse”. (ii) A collapsed fork can also be rescued by a converging RF. This strategy is efficient due to an excess of potential replication origins (dormant origins) that can be activated upon replication stress^5^. However, some fragile sites have a paucity of origin activity over megabase distances^6^. These regions are particularly sensitive to replication problems, likely because two converging forks that are forced to travel long distances are more likely to both collapse and there are no dormant origins present in the intervening region. (iii) To overcome the consequences of two converging forks collapsing, or the collapse of single forks in regions of unidirectional replication such as telomeres, cells have developed mechanisms to restart a collapsed RF by homologous recombination (HR)^7^.

The major pathway of HR-mediated fork restart requires Rad51 and the classical HR mediators. Restart can occur from a one-ended DNA double strand break (DSB), such as would occur if the RF encounters a nick^8^, or from a resected replication fork, likely following fork reversal that generates a 4-way DNA junction known as a “chicken-foot”^9^. To explore the latter mechanism of HR-restart of collapsed RFs we developed a model system in fission yeast using a unidirectional replication fork barrier, *RTS1*{Lambert et al., 2005, #90478}. Using this system, we and others have demonstrated that HR-dependent restart occurs rapidly (∼18 minutes^10^), does not require a DSB intermediate^11^ and likely involves an initial fork reversal step that allows regulation of resection via Ku binding and subsequent displacement by the Mre11-Rad50-Nbs1 complex^12^. We and others have also shown that the restart of arrested forks is associated with various types of genetic instability: ectopic recombination events can restart the fork at the wrong locus causing rearrangements such as translocations^11^, while forks restarted at the correct locus are intrinsically more error prone than a canonical fork. The errors intrinsic to the restarted replication machine include increased replication slippage^13^, recombination between direct repeats^14^ and the coupled formation of dicentric and acentric isochromosomes at inverted repeats^15^. The increase in error frequency implies that HR-restarted RFs are different from the canonical RF. Indeed, we found that, unlike canonical replication (where polymerase Pol ε synthesises the leading strand and Pol δ the lagging strand), both the leading and lagging strands are synthesised by Pol δ despite the fact that replication remains semi-conservative^10^ and does not therefore progress as a migrating D-loop as is seen during break-induced replication^16^.

One limitation of current assays to characterise fork restart at *RTS1* is the inability to track replication dynamics around the site of the restarted fork. Thus, many conclusions are drawn from assays that measure ectopic recombination, the mutagenic potential of the restarted RF or which quantify specific DNA structures using 2D-gels. Here we combine polymerase usage sequencing (Pu-seq:^17^) with a Monte Carlo model of *S. pombe* DNA replication^18^ to complement these assays. Pu-seq relies on mutant replicative polymerases that introduce an increased frequency of ribonucleotides during DNA synthesis^19^. By mapping the strand-specific location of ribonucleotides across the genome and comparing the profiles of strains mutated in the different replicative polymerases, Pu-seq tracks polymerase usage across the genome. Pu-seq thus allows the direct visualisation of restart *in vivo* and is complementary to the above mentioned indirect indicators such as mutagenic outcomes. By modelling Pu-seq derived replication dynamics and comparing this with experimental data we are able to gain significant new insights into the process of HR-dependent replication restart.

Consistent with our previous work^10^, we confirm that Pol δ synthesises both the leading and lagging strands (referred to as δ/δ configuration) and demonstrate that the HR-restarted replication machine does not “mature” into an ε/δ configuration over ∼30 kb, but continues in the δ/δ configuration until it is terminated upon encountering a converging fork. We also provide direct evidence that Pol α does not participate significantly in the DNA synthesis mediated by the HR-restarted RF. This suggests that the lagging strand template remains as a gap that is filled-in following fork merger.

During fork restart the lagging strand is degraded and, following Rad51-dependent strand invasion, the leading strand acts as the template to initiate replication. To explore the stability of the leading stand we manipulated the timing of restart and the extent of resection behind the fork arrested at *RTS1* by deleting *pku70*^12^. We confirm that extending resection of the lagging strand impacts on the timing of HR-restart but does not influence site of HR-restarted fork initiation. This suggests that the leading strand 3’ end remains stable.

## Results

Replication forks are efficiently arrested at *RTS1*, a programmed unidirectional replication fork barrier of ∼850bp DNA derived from the mating type locus of fission yeast^20^. *RTS1* itself is not intrinsically difficult to replicate and is only active as a fork barrier when bound by Rtf1{Eydmann et al., 2008, #4322}. To maximise the distance that a restarted fork travels we tested four locations, introducing the *RTS1* sequence immediately adjacent to an efficient early firing origin that boarders a transition zone between early and late replicating regions^17^ and performed Pu-seq on each in the presence of *RTS1* barrier activity (Figure S1). From the resulting Pu-seq traces we chose a locus on chromosome II, which showed the highest extent of HR-restarted replication over distance. At this locus (Figure 1A), when the left-to-right RF emerges from the early origin and reaches *RTS1*, it is arrested and subsequently restarted by HR. During this time, the distant (approximately 30 kb away) late origin is activated, but the difference in timing of origin activation and the time taken for fork movement means that the converging right-to-left moving fork does not reach the *RTS1* sequence and the HR-restarted RF travels left-to-right for approximately 5-15 kb, until it terminates with the convergent right-left fork.

**Figure 1.**
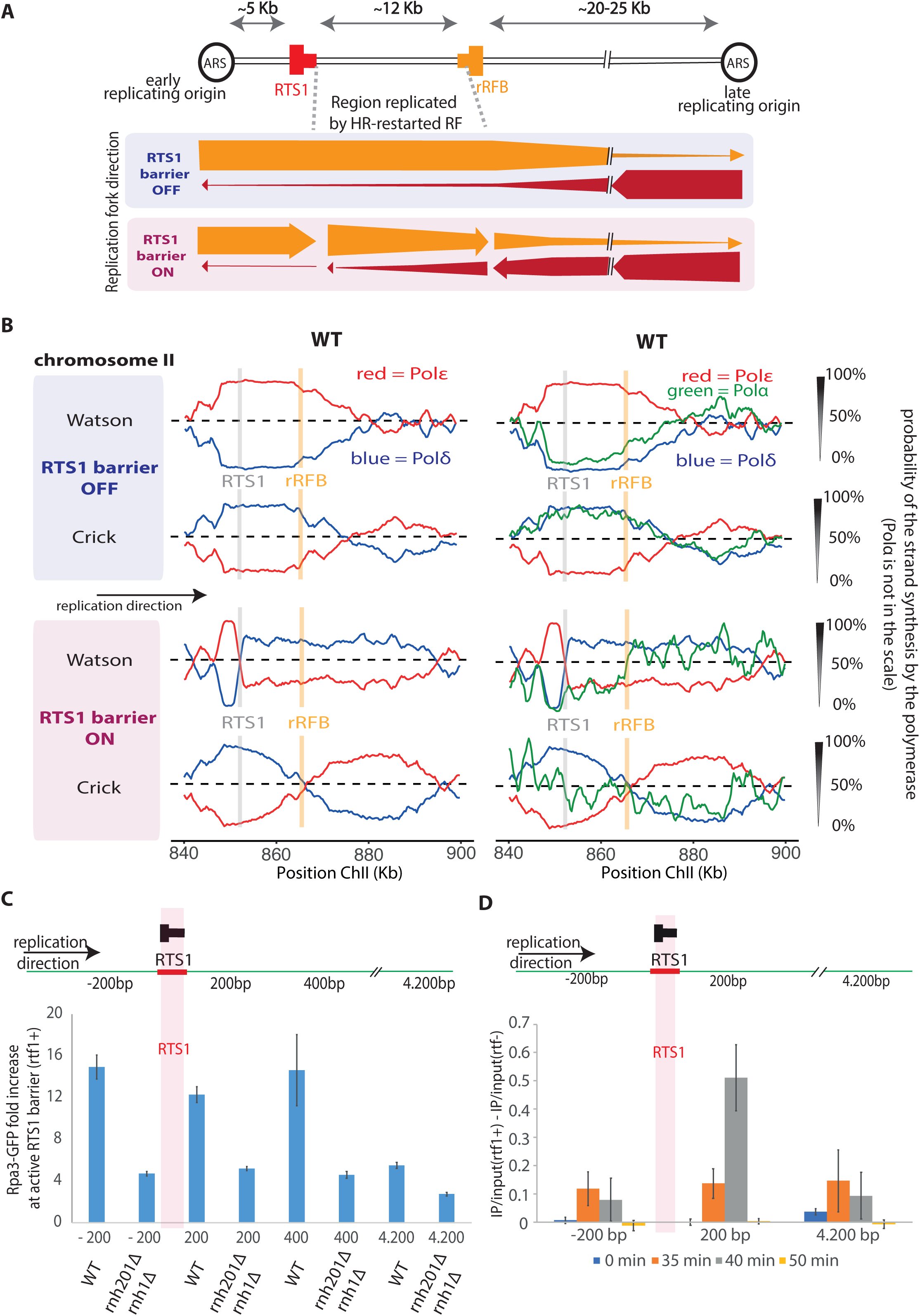
Polymerase usage following HR-restart. **A**. Schematic of the *RTS1-rRFB* locus on chromosome II. The positions of the directional *RTS1* and rRFB barriers are shown as red and orange, respectively. The thick bar represents the directionality of fork arrest. ARS: autonomously replicating sequence. The direction (see panel B) of unperturbed and perturbed replication at this locus is indicated by the thickness of arrows underneath. **B**. Pu-seq traces of the ChrII locus. Top two traces: *RTS1* barrier activity off (*rtf1-d*). Bottom two traces: *RTS1* barrier activity on (*rtf1*^+^). Left panels: the usage of Pol δ (blue) and Pol ε (red) are shown on the Watson and Crick strands. Note the switch from Pol ε to Pol δ on the Watson strand at the *RTS1* site is indicative of a change in polymerase usage on the leading strand when *RTS1* barrier activity is on. Right: The same traces overlaid with data for Pol α (green). **C**. Chromatin immunoprecipitation of Rpa3-GFP at the indicated positions (relative to base 1 of the *RTS1* sequence) in unsynchronised cells with either *rnh1*^+^ *rnh201*^+^ (WT) or *rnh1-d rnh201-d* backgrounds. Data presented are relative to the *ade6*^+^ control locus. n = 3 biological repeats. Error bars: standard deviation of the mean. **D**. Chromatin immunoprecipitation using the S9.6 antibody in wild type cells synchronised in G2 and released into S phase. n = 3 biological repeats. Error bars: standard deviation of the mean.

To further delay the converging fork, we next inserted 10x copies of the Ter2-Ter3 sequence^21^ from the *S. pombe* ribosomal DNA replication fork barrier (rRFB) 12 kb downstream of *RTS1*, between *RTS1* and the late firing origin (Figure 1A). At the rRFB barrier, RFs do not collapse^22^, likely because the rRFB binding proteins activate the fork protection complex^23^, and the RF is only transiently paused and does not undergo HR restart.

To establish the extent of the delay caused by the rDNA barrier, we first inserted the 10x Ter2-Ter3 close to a strong origin (on ChrI) and used Pu-seq to monitor fork direction across the locus. We then implemented a recently described Monte Carlo model of *S. pombe* DNA replication^18^ that we have modified to account for programmed replication barriers (see Materials and Methods). Using this program, we modelled various delay scenarios. The best match to the experimental data was ∼6-minute delay for the rRFB construct (Figure S2).

Using our optimised construct for Pu-seq we observe that the majority of the 12 kb between the *RTS1* and rRFB is replicated by the HR-restarted fork when the barrier is active (Figure 1A,B). In previous work using qPCR immediately upstream and downstream of the RTS sequence we have estimated the delay time at the *RTS1* barrier to be ∼18 minutes^10^. By applying our modified model, we show that the best fit for the experimental data is a delay at *RTS1* of 11 minutes (Figure S3). It should be noted, however, that we assume in the model that replication speed following restart is equivalent to canonical replication (we used 1.8 kb/minute). If this assumption was incorrect, the optimal delay time would change. For example, if we model HR restarted fork speed at 3kb, the optimal delay time from the model is 17.5 min; see discussion.

### Polymerase alpha usage during HR-restarted replication

In our original description of Pu-seq^17^ we used two strains for each experiment: one containing a mutation in the gene encoding the catalytic subunit of Pol δ (*cdc6-L591G* or *cdc6-L591M*) and one containing an equivalent mutation in the gene encoding the catalytic subunit of Pol ε (*cdc20-M630F*). To monitor the contribution of polymerase α, we included an additional strain containing an equivalent mutation in the gene encoding the catalytic subunit of Pol α (*swi7-L850F*). Previously we suggested that Pol α contributed less to replication of the lagging strand by HR-restarted forks than for canonical forks because Pol α dependent mutagenesis was reduced approximately 6-fold when forks were replicated by HR-restarted replication^10^. From the Pu-seq traces (Figure 1B; also see Figure S4) we now confirm and extend this: Pol α usage is significantly reduced during synthesis of the lagging strand when replication is performed by an HR-restarted fork.

This observation is consistent with a model whereby the HR-restarted fork synthesises the complete leading strand without coupled Okazaki fragment synthesis on the lagging strand. Presumably, upon termination, the “lagging strand” is then replicated by Pol δ from the 3’ end derived from the converging forks leading strand. This would result in RPA-bound ssDNA accumulating significantly in the region and the possibility that RNA:DNA hybrids may also accumulate^24^. To establish if this was the case, we examined the accumulation of RPA by ChIP at several sites around the locus in asynchronous *rtf1*^+^ cells (∼10% of which will be in S phase) (Figure 1C). We observed significant RPA accumulation at that part of the locus replicated by an HR-restarted fork when normalised against a locus (*ade6*) that was replicated by canonical forks. This was attenuated in an *rnh201-d rnh1-d* double mutant background, suggesting competition between RPA and RNA:DNA hybrids, as previously reported at resected double strand breaks^24^. Using an antibody with specificity to RNA:DNA hybrids we also saw evidence for the accumulation of RNA:DNA hybrids in S phase (Figure 1D).

### HR-restarted replication forks do not mature to an ε/δ configuration

We had previously reported that HR-dependent replication is semi-conservative and proceeds in a manner by which Pol δ synthesises both the leading and lagging strand (δ/δ configuration). We did not see any evidence that HR-dependent replication reverted to a more canonical ε/δ configuration over 2-3 kb^10^. The data in Figure 1B extend this: the fact that there is no increased Pol ε usage between the *RTS1* and rRFB sites (∼12 kb) on the leading (Watson) strand implies there is no maturation of the HR-restarted replication machine to the canonical distribution of polymerase labour

To explore this further we designed an additional construct where a second *RTS1* sequence was integrated in the opposing orientation to the first, and close to the late origin (Figure 2A). This second *RTS1* sequence was intended to arrest the right-to-left fork emerging from the late origin and promote its HR-restart. The rationale was to isolate an origin-free region of ∼30 kb and force this to be replicated by HR-restarted forks. If HR-restarted forks do transition from δ/δ configuration to ε/δ, then we would predict significant evidence of Pol ε usage in the Pu-seq traces. As is evident (Figure 2B) we see no significant Pol ε usage on either strand across >30 kb, commensurate with HR-restarted forks replicating consistently in the δ/δ configuration.

**Figure 2.**
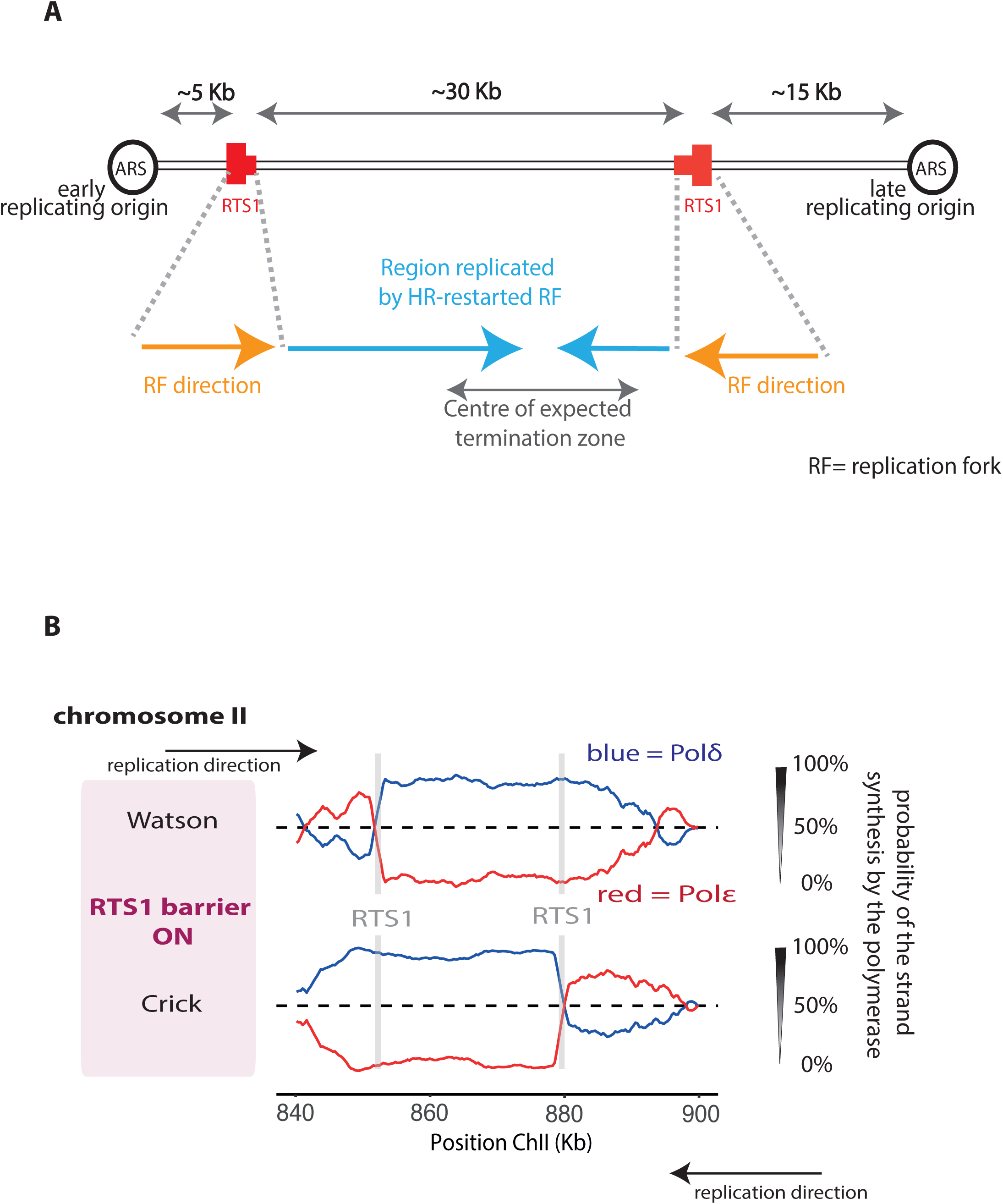
HR-restarted replication does not mature from the δ/δ to ε/δ configuration. **A**. Schematic of the inverted *RTS1* locus on chromosome II. The positions of the directional *RTS1* barriers are shown in red, with the thick bar indicating the directionality of fork arrest. **B**. Pu-seq traces from the modified locus when the barrier is active (adh-*rtf1*^+^). Note: the transition from Pol ε (red) to Pol δ (blue) on the Watson strand at the left-side *RTS1* site and the transition from Pol ε to Pol δ on the Crick strand at right-side *RTS1* site.

### HR-restarted forks are insensitive to the *RTS1* barrier

In previous work using 2D gels we estimated that replication arrest at *RTS1* was >90% efficient^11^. However, in the traces shown in Figure 1B only ∼70% of forks appeared to switch replication from an ε/δ to a δ/δ configuration. We reasoned that this may reflect the fact that, in these experiments, we are relying on endogenous *rtf1*^+^ levels while in previous work we regulated the expression of *rtf1*^+^ through use of an inducible *nmt1* promoter. To test the possibility that the level of *rtf1*^+^ transcription influences the efficiency of arrest at *RTS1*, we replaced the endogenous *rtf1*^+^ promoter with the *adh1*^+^ promotor (adh-*rtf1*^+^) and performed an additional Pu-seq experiment (Figure 3A). When compared to *rtf1*^+^, the adh*-rtf1*^+^ traces indeed showed an increased transition from the ε/δ to the δ/δ configuration. Using our model of DNA replication, we estimated that the efficiency of *RTS1* barrier in arresting a canonical fork in the strain with endogenous *rtf1*^+^ reached ∼70% but in adh*-rtf1*^+^, the blocking capacity of *RTS1* increased to 90% (Figure S5). We thus conclude that the *RTS1* barrier is indeed highly efficient when Rtf1 is not limiting.

**Figure 3.**
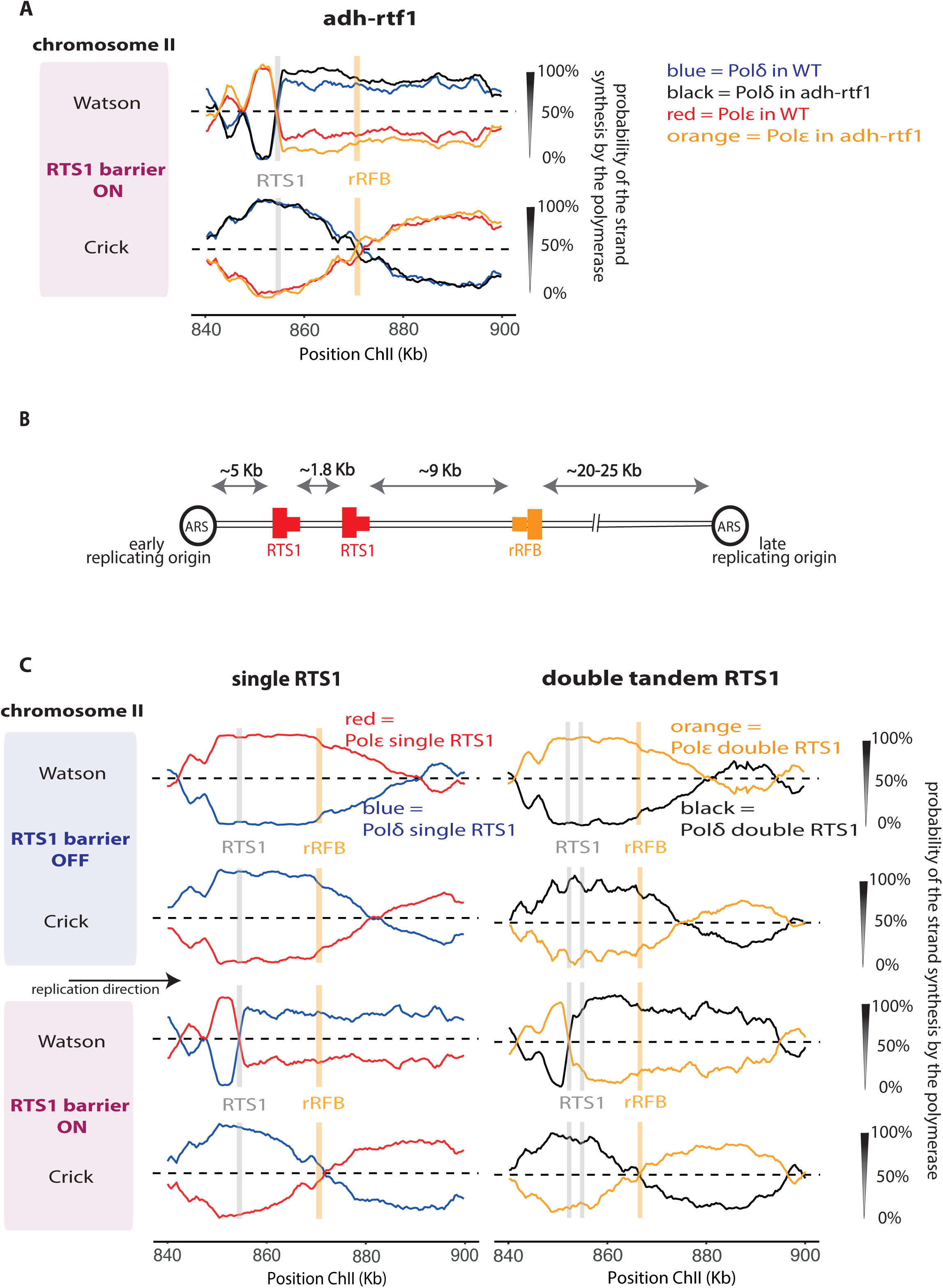
HR-restarted forks are insensitive to the *RTS1* barrier. **A**. Pu-seq traces for the *RTS1-rRFB* locus are overlaid for two sets of strains: *rtf1*^+^ (WT), which has endogenous *rtf1* expression (Pol δ: blue, Pol: ε red) and adh-*rtf1*^+^, where expression is under control of the *adh1* promoter (Pol δ: black, Pol: ε orange). **B**. Schematic of the tandem *RTS1-rRFB* locus on chromosome II. The positions of the directional *RTS1* and rRFB barriers are shown as red and orange, respectively with the thick bar indicating the directionality of fork arrest. ARS: autonomously replicating sequence. **C**. Pu-seq traces for the tandem *RTS1-rRFB* locus which contains two copies of the *RTS1* barrier that are both orientated to arrest left-right forks (right panel) compared to the original *RTS1-rRFB* locus (left panel). Top panel: barrier off. Bottom panel: barrier on.

The fact that the *rtf1*^+^ experiment only arrests ∼70% of forks prompted us to address a longstanding question: does the HR-restarted fork also get arrested at an *RTF1* barrier, or is it intrinsically resistant to this particular barrier? If the latter were the case, it would explain why the HR-restarted fork can overcome the initial barrier, which stopped the canonical fork. We therefore modified the locus to include a second copy of *RTS1* approximately 1.8 kb (Figure 3B) downstream and performed additional Pu-seq experiments with and without the presence of *rtf1*^+^ (Figure 3C). In the absence of fork arrest the traces of polymerase usage are as expected, with the vast majority of forks travelling left-right. In the presence of *rtf1*^+^, again as expected, ∼70% of left-right forks arrest at the first *RTS1* sequence and restart in a δ/δ configuration. This is indicated by a sharp transition in Pol δ usage on the Watson strand at the first *RTS1* site. At the second *RTS1* sequence, the majority of the remaining 30% of canonical forks that were not arrested at the first *RTS1* barrier arrest and restart at the second *RTS1* site. This is indicated by the additional Pol δ usage inflection on the Watson strand at the second *RTS1* site.

We can thus evaluate if the 70% of HR-restarted forks that were arrested at the first barrier are significantly delayed as they pass through the second barrier by comparing the position of terminations (fork merger events). This can be seen from the position and gradient of Pol ε usage on the Crick strand. In this context Figure 3C demonstrates two things: first, activating either one or two barriers significantly shifts the position of the termination events to the left, commensurate with a delay to left-right replication forks. Second, when the barrier(s) are active, termination positions and frequencies do not change significantly between the situations where there is either one or two barriers. Thus, we can conclude that the second *RTS1* barrier does not significantly delay the progress of HR-restarted replication forks. Again, we can apply our model to the scenario, applying different delays to forks restarted at the first barrier when they pass through the second barrier (Figure S6). The best fit to the experimental data is no delay of HR-restarted forks at the second barrier. We thus propose that HR-restarted forks are unaffected by the *RTS1* barrier.

### Altering resection of the lagging strand upstream of the arrested fork

A role for Ku at *RTS1* arrested forks was recently reported^12^. By measuring ssDNA formation using a restriction enzyme protection assay it was determined that, in the absence of *pku70*, the distance to which resection progressed behind the arrested fork was increased by a factor of greater than 2, to beyond 1800 bp. Unexpectedly, using quantitative microscopy and chromatin immunoprecipitation in the region between 0 and 600 bases behind the fork, RPA and Rad51 association were shown to be decreased approximately 2 fold. Using a genetic assay for replication slippage to monitor replication by non-canonical forks this was correlated with a decrease in HR-restarted forks downstream of the *RTS1* barrier. Thus, it was proposed that replication restart by HR was delayed in the absence of Ku, potentially because Ku is required to impose regulated MRN/CtIP- and ExoI-dependent two-step resection at arrested forks.

To address the consequences of Ku loss on replication dynamics we introduced a *pku70* deletion mutation into our Pu-seq strains. We also modified the Pu-seq protocol by introducing a *rnh201-RED* mutation (RED; ribonuclease excision defective) instead of the *rnh201* deletion. RNase H2 can remove single ribose bases in DNA through initiating the ribonucleotide excision repair (RER) pathway^25^. It can also degrade DNA:RNA hybrids. Preventing RER is essential to the methodology of Pu-seq, but the loss of any DNA:RNA hybrid activity is not required and has the potential to impact on some aspects of replication. A separation of function mutation, *RNH2-RED* has been described and characterised in *S. cerevisiae*^26^. We created a mutated *rnh201* resulting in the equivalent change (Rnh201-P75D:Y245A) and characterised the strain genetically in *S. pombe* to demonstrate successful separation of function (Figure S7).

We performed Pu-seq in the presence or absence of *pku70*^+^ in either the *rnh201-d* or *rnh201-RED* backgrounds. Genome-wide we observed no significant difference between *rnh201-d* and *rnh201-RED*. We next plotted the traces for both “arrest off” and “arrest on” conditions to visualise the *RTS1* locus (Figure 4A). Following arrest and restart we observed that the *rnh201-RED* mutation resulted in no difference to the traces from either the *pku70*^+^ or the *pku70-d* backgrounds. We thus conclude that the specific allele of *rnh201* that we use does not influence the experiment.

**Figure 4.**
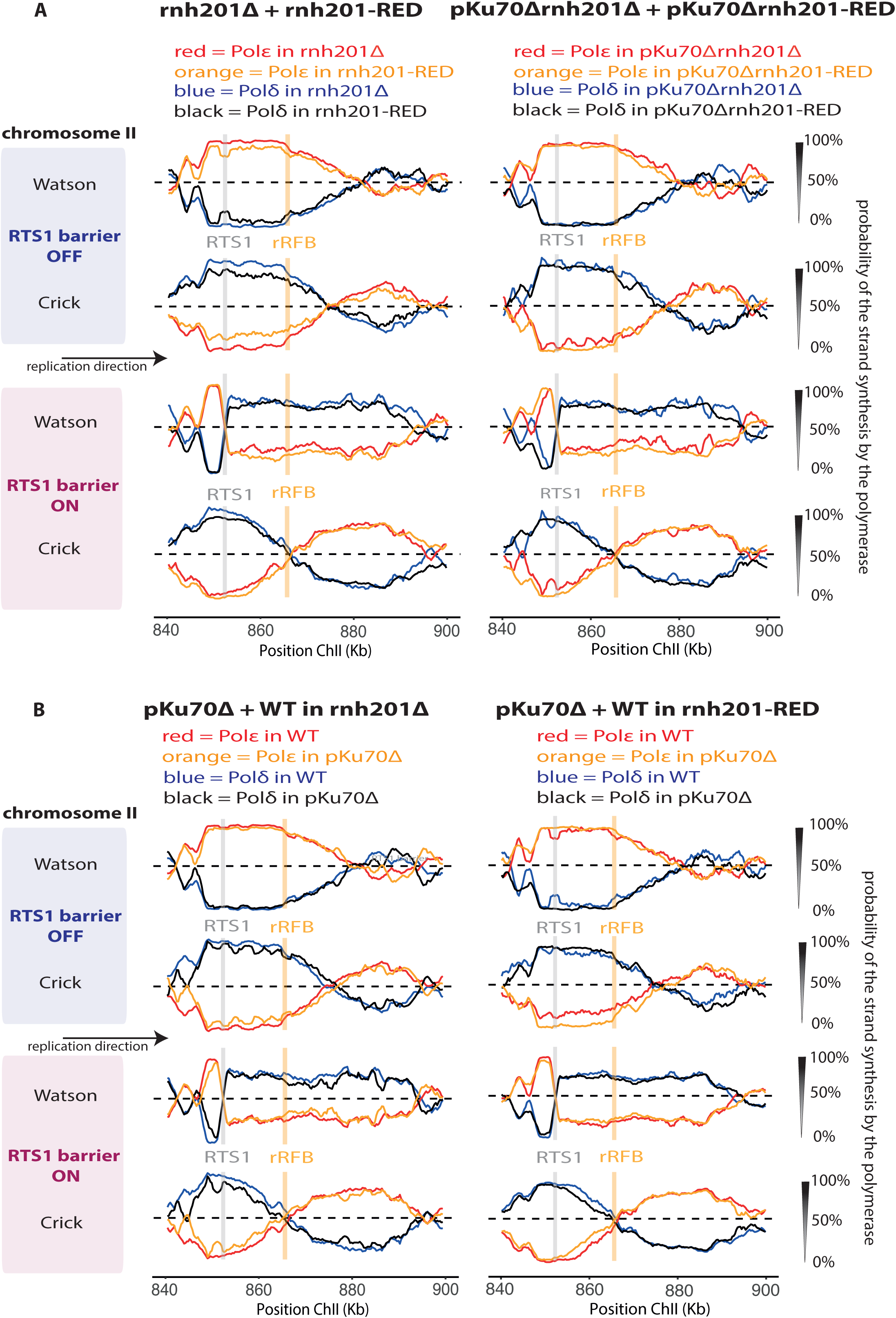
Loss of *pku70*^+^ results does not destabilise the leading stand primer. **A**. Overlays of Pu-seq traces of the *RTS1-rRFB* locus either with or without arrest and in the presence and absence of *pku70*^+^. Left panel: in the *rnh201-d* background. Right panel: in the *rnh201-RED* background. **B**. For the purpose of comparison, overlays of the Pu-seq traces for *rnh201-d* and *rnh201-RED* backgrounds. Left panel: in *pku70*^*+*^. Right panel: in *pku70-d*.

The presence or absence of Pku70 does not cause a major change in the Pu-seq profiles after fork arrest. However, a careful examination of the data (Figure 4B) suggests a slight shift in the overall position of termination events (evidenced by the position and slope of Pol ε usage on the bottom strand) that would be consistent with an increased time for replication restart. The apparent delays are visible in both the *rnh2-d* and *rnh2-RED* backgrounds. Using our model of DNA replication, we can estimate that the extent of the delay to HR-restart of forks arrested at *RTS1* in the absence of Pku70 as 14 minutes, which is an increase of 3 minutes (∼27%) when compared to *pku70*^+^ background. (Figure S8). These data confirm that Ku loss has a direct effect on the dynamics of replication fork restart^12^.

One potential explanation for the altered dynamics of replication that we observe in the *pku70* mutant is that HR-restart occurs further upstream (left) of the original fork arrest site (see discussion): the increase in resection reported to occur behind the fork in the *pku70-d* mutant could affect the stability of the leading strand that provides the invading strand during HR and primes HR-restarted replication. If the leading strand were degraded, we would expect to observe strand invasion and thus new δ/δ configured DNA synthesis occurring further upstream of the *RTS1* sequence. However, in the presence or absence of Pku70, the transition from Pol ε to Pol δ on the Crick strand, which marks the switch of polymerases, is unchanged (Figure 4). Thus, we conclude that the leading strand remains stable, despite the increased resection and the delay to restart.

## Discussion

The increased genomic instability that is caused by replication stress when cancer cells are driven into uncoordinated replication by oncogene activation may, in part, be caused by the HR-mediated replication restart. To further understand this driver of genetic stability we have applied polymerase usage sequencing^19^ to characterise the *RTS1* model replication restart system that had previously been established in *S. pombe*. As expected, our data confirmed the previous conclusions that, when replication is restarted by HR, Pol δ is used to replicate both the leading and lagging strands^10^. We have also been able to extend several previous observations: we demonstrate that HR-restarted replication forks do not “mature” into the more canonical ε/δ conformation over a distance ∼30 kb and that Pol α is not used at significant levels during HR-restarted replication. This latter observation suggests that the leading and lagging strands are replicated consecutively, rather than being coupled as they are during canonical replication. In support of this we demonstrate that both RPA and potential DNA:RNA hybrids are strongly enriched at the locus that is replicated by HR-restarted replication.

### The dynamics of replication restart at *RTS1*

When a replication fork collapses and is unable to resume canonical replication, it can be rescued by a converging fork. However, previous work has shown that a single *RTS1* barrier frequently results in HR-dependent replication fork restart even though, in the vast majority of cells, a converging fork would be expected to arrive^27^. This implies that HR-dependent restart occurs irrespective of the status of the locus, but is instead governed by the time it takes to assemble the appropriate machinery. Using qPCR upstream and downstream of *RTS1* we have previously estimated that restart occurs after a delay of ∼18 minutes in the vast majority of cells^10^. A separate study demonstrated that HR protein foci begin to appear 10 minutes after the start of S phase, peak in numbers some 30 minutes later and that individual foci can remain present for >30 minutes^27^. To reconcile these two data sets, we have speculated that HR-mediated restart occurs in most cells after a delay of ∼18 minutes and that HR proteins remain associated with the locus during HR-restarted replication progress (and possibly during fork convergence and termination with the incoming canonical fork).

The utility of combining a physical profile of replication dynamics with a mathematical model of DNA replication that can incorporate complementary genetic and physical data is exemplified by our consideration of the duration of replication arrest at *RTS1*. By extending a previously reported Monte Carlo simulation of *S. pombe* DNA replication^18^ to include the ability to modify replication speed in a time and location dependent manner (see Materials and Methods) we observed that the best fit of the model to the experimental data (given by the Euclidean distance) is a delay of 11 minutes at *RTS1*, before replication is restarted by HR. This estimate is quite distinct from the observed experimental data obtained by qPCR of 18 minutes^10^. A potential explanation for this discrepancy is that the timing estimate assumes that the speed of HR-restarted replication is equivalent to the speed of the canonical fork (which we modelled at 1.8 kb/min). However, in *S. cerevisiae*, it has been estimated that break-induced replication, which proceeds via a migrating D loop, progresses at between 3 and 4 kb a minute^28^. If we increase the speed of the HR-restarted fork to 3 kb in the model, then our estimate of the delay to DNA replication at *RTS1* is extended to 17.5 minutes. Thus, based on the qPCR evidence of replication time upstream and downstream of *RTS1* varying by 18 minutes when the barrier is active, we propose that HR-dependent replication restarts relatively synchronously and proceeds approximately 50% faster than canonical replication.

### The HR-restarted fork is insensitive to the *RTS1*/Rtf1 barrier

When a fork is restarted by HR following arrest at the *RTS1* barrier, it clearly is able to progress through the barrier. We have previously speculated that the restarted replication apparatus is distinct from the canonical replisome and that the HR-dependent apparatus is not sensitive to the barrier^15^. Here we have tested this by following the dynamics of HR-restarted forks as they pass through a second copy of *RTS1*. Visual examination of the data suggests no appreciable evidence of a delay to the HR-restart replication imposed by the second barrier. By modelling a variety of scenarios whereby HR-restarted forks are delayed between 0 and 30 minutes at the second barrier, we demonstrate that the best fit of the experimental data to the model is 0 minutes delay. This demonstrates that HR-restarted replication is insensitive to at least this obstacle, which efficiently arrests canonical forks. Thus, the distinct nature of the restarted apparatus may provide cells with an alternative replication machine that is able to pass some barriers more effectively. Clearly, this comes at a cost - the HR restarted fork is highly error prone and, since we also provide evidence that leading and lagging strand synthesis are not coupled, significant quantities of ssDNA will be present that would be sensitive to oxidative stress^29^. The increased errors that occur when replication is restarted by HR may be a driving force behind the evolution of the MCM8/9 helicase, which is proposed to function in metazoan cells in HR-dependent replication^30^

### Replication restart is delayed in Ku mutants

Having established a system to track replication dynamics we were interested to explore the effects of deregulating resection by deletion of *pku70*. A recent study demonstrated that loss of Ku caused the loss of Rad50- and Ctp1-dependent two-stage resection regulation and resulted in extended tracts of ssDNA that stretch several kilobases behind the arrested fork^12^. By measuring the error rate in a replication slippage assay, the authors suggested that, because the error rate decreased by ∼50% and this could be rescued by delaying the incoming fork, this could be interpreted as a delay to HR-dependent restart. The most prosaic expectation would predict that the loss of the requirement that Ku70 to be removed from a reversed fork by MRN would actually lead to faster restart. We considered an alternative explanation for the delay: that 3’ end which primes the strand invasion event is unstable and, during the extended 5’-3’ exonuclease activity, it is shortened. This would result in replication restart occurring further upstream from the initial arrest site. Even if the restart occurred at the normal time (i.e. after approximately 18 minus), this would manifest as a delay, since the HR-restarted fork would have further to travel and thus an increased proportion of the locus would be replicated by the converging canonical fork.

Using our Pu-seq assay we were able to demonstrate that the transition from Pol ε to Pol δ occurs at the same position in *pku70*^+^ and *pku70*-*d* backgrounds. Since the position of this transition is caused by the switch from Pol ε during canonical replication to Pol δ when replication is restarted by HR, this demonstrates that the 3’ end remains stable even when resection is deregulated by loss of *pku70* and restart occurs at the same place irrespective of Ku status. By modelling the kinetics of replication, we were able to confirm that loss of Pku70 indeed resulted in an additional delay to replication restart of 3 minutes (an additional 27% compared to *pku70*^+^). This manifests as a shift in the positions of termination of the HR-restarted forks with converging canonical forks.

It could be asked why a 27% increase in the time taken to restart DNA replication in the absence of Ku changes the mutation frequency immediately downstream of the *RTS1* barrier by a factor of two^12^. It should be noted that these genetic experiments were performed using the *RTS1* barrier integrated at the *ura4* locus on chromosome III. At this locus, as opposed to our optimised ChrII locus, there are more forks converging on the arrest site from active downstream origins and thus a modest delay to restart would be expected to have a significantly more pronounced effect on the percentage of DNA downstream of the barrier that is replicated by an HR-restarted (error prone) fork as opposed to a converging canonical fork. It remains unclear why there is a delay to restart in the *pku70-d* background, although it is proposed to be linked to the reduction of RPA and Rad51 observed by microscopy^12^.

## Acknowledgments

This work was funded by the Wellcome Trust. Grant number: 110047/Z/15/Z.

## Author contributions

KN, EC-F and AMC conceived and designed the study. KN, ATW, AB and IM acquired the data, KN, EC-F and AMC interpreted the data. KN and AMC drafted the manuscript. All authors edited the manuscript.

## Data availability

The datasets generated during the current study are available in the NCBI GEO repository: [https://www.ncbi.nlm.nih.gov/geo/query/acc.cgi?acc=GSE153731].

## Code availability

The code for the model used is available at: [https://github.com/ecam85/fork_barrier]

## Supplementary Figure legends

**Figure S1.**
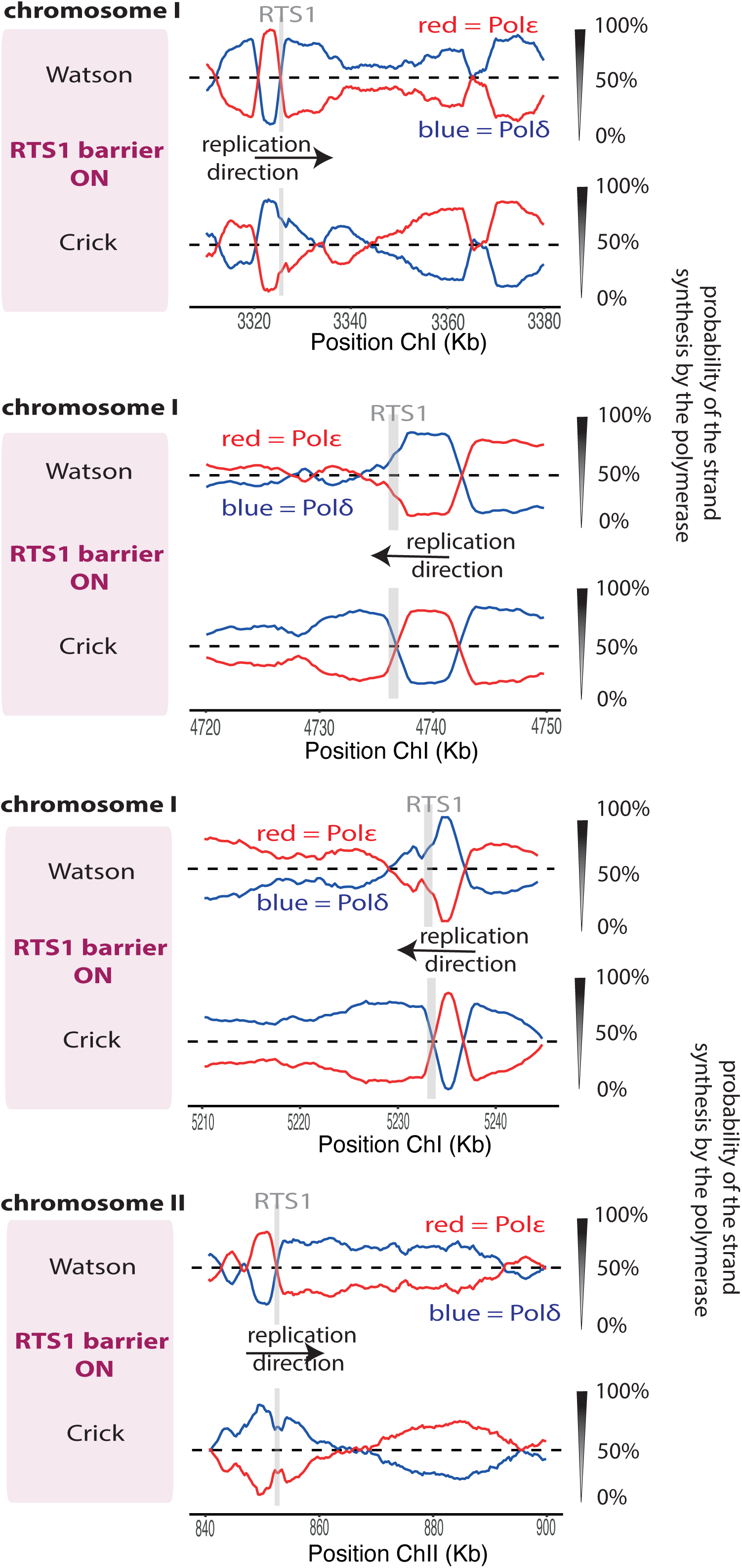
Polymerase usage following HR-restart at 4 different loci. Pu-seq traces at three loci on ChrI and one locus on ChrII where the *RTS1* barrier has been integrated close to a strong origin in an early replicating region of the chromosome that boarders a late replicating region. *RTS1* barrier activity on (*rtf1*^+^). The usage of Pol δ (blue) and Pol ε (red) are shown on the Watson and Crick strands. Note the switch from Pol ε to Pol δ at the *RTS1* site that is indicative of a change in polymerase usage from Pol ε to Pol δ on the leading strand when *RTS1* barrier activity is on and replication is restarted by HR.

**Figure S2.**
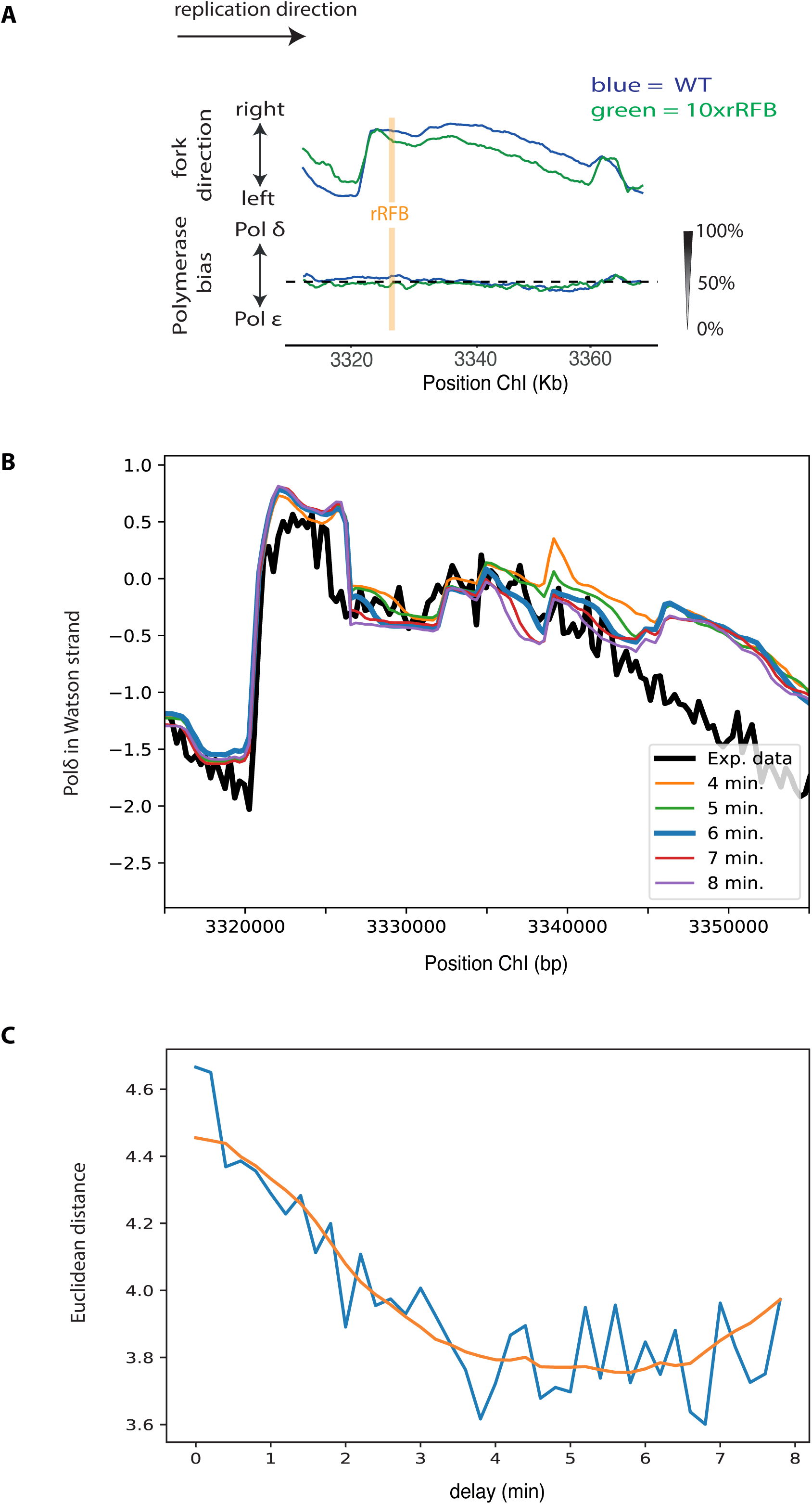
Calculating the delay to replication at rRFB barrier. Pu-seq traces of a locus on ChrI with and without the integration of 10xTer2-Ter3 sequence (rRFB barrier). **A**. Top Panel: Fork direction calculated from the Pu-seq traces (see^17^). When the rRFB barrier construct is integrated, there is a decrease in the proportion of left-right moving forks downstream of the barrier. This is caused by a delay to left-right moving forks, and a concomitant increase in right-left moving forks. Bottom panel. Polymerase delta usage averaged across both strands is shown, demonstrating that (unlike for *RTS1*) forks only pause and are not restarted by HR.**B**. Selected outputs (4 - 8 mins, 1 minute intervals) of the Monte Carlo model for the indicated delays to the left-right replication forks at rRFB are compared to the experimental data. **C**. Error between the experimental data and the model output for a delay between 0 and 8 minutes (15 second intervals) measured by the Euclidean distance between the two signals; orange curve is a smoothed representation of the errors using a Savitzky-Golay filter.

**Figure S3.**
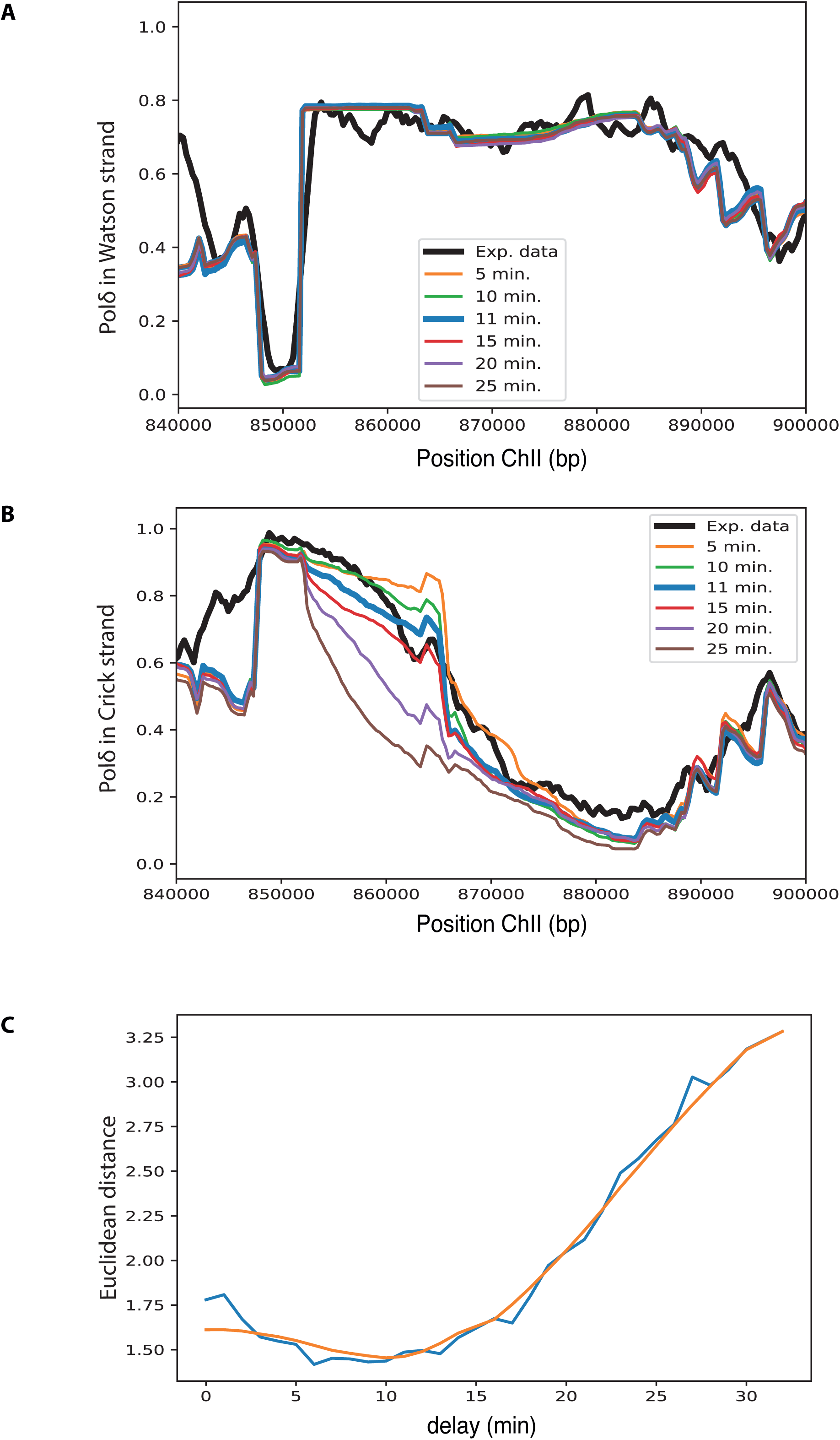
Calculating the delay at *RTS1* barrier. Using the replication model, different delay times of replication at the *RTS1* barrier before HR-dependent restart (between 0 and 30 minutes, 1 minute steps) were fitted and compared to the experimental data. The best fit is an 11 minute delay at the barrier. **A**. Selected outputs from the model for usage of Pol δ on Watson strand are compared to the experimental data. Note: both when replicated by left-right HR restarted forks, or when replicated by right-left converging canonical forks, the Watson strand is replicated by Pol δ. **B**. Selected outputs from the model for usage of Pol δ on Crick strand are compared to the experimental data. Note: A decrease in Pol δ usage on the Crick strand reflects replication by Pol ε from right-left converging forks. Thus, because an increased delay results in more replication by converging canonical forks, this is reflected by a decrease in Pol δ usage. **C**. Error between the acquired data and the model output measured by the Euclidean distance between the two signals; orange curve is a smoothed representation of the errors using a Savitzky-Golay filter.

**Figure S4.**
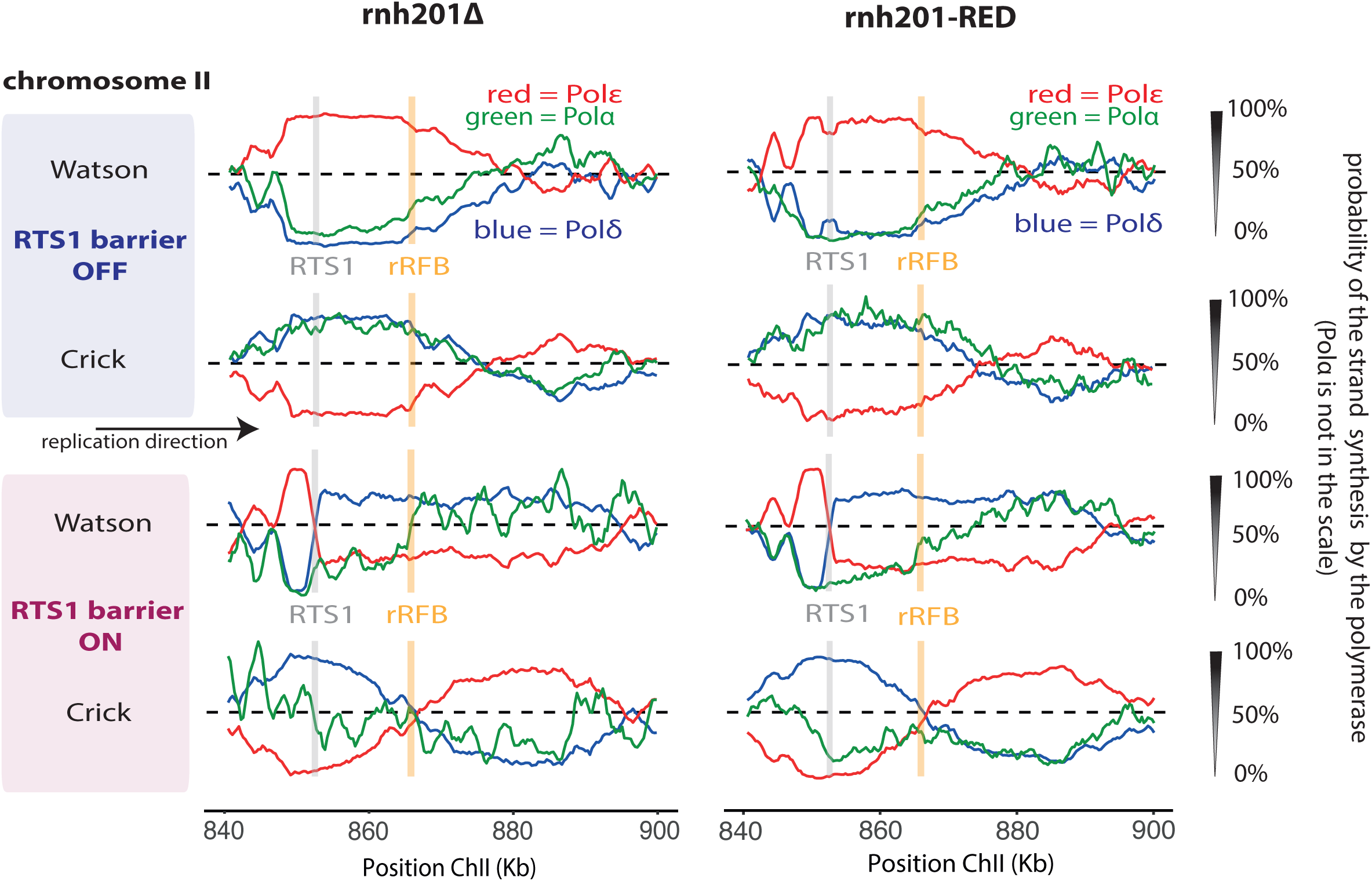
Comparative polymerase usage following HR-restart between *rnh201-d* and *rnh201-RED*. Pu-seq traces of the *RTS1-rRFB* locus on ChrII. Top two traces: *RTS1* barrier activity off (*rts1-d*). Bottom two traces: *RTS1* barrier activity on (*rtf1*^+^). The usage of Pol δ (blue), Pol ε (red), and Pol α (green) are shown on the Watson and Crick strands in *rnh201-d* and *rnh201-RED*.

**Figure S5.**
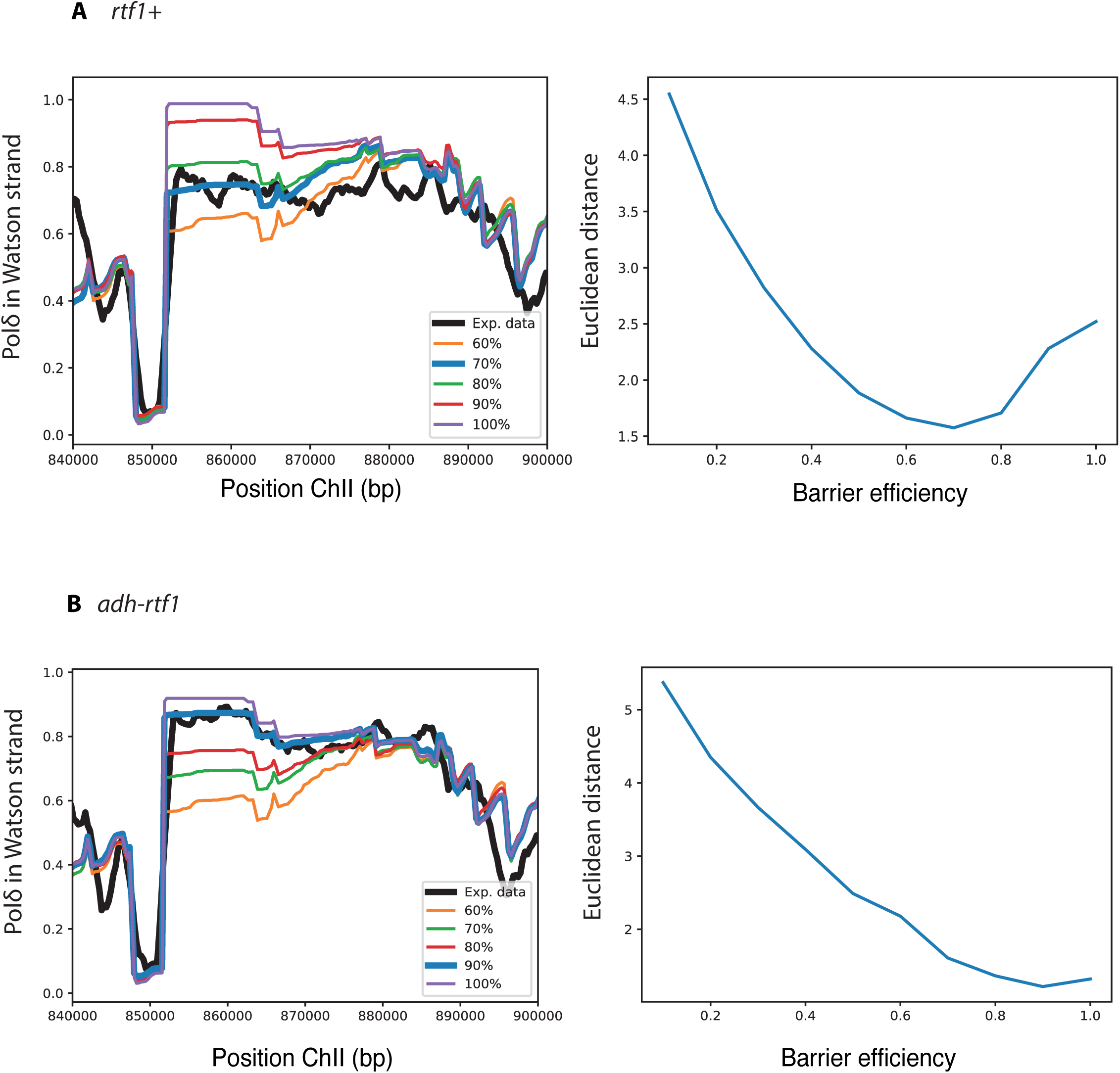
Efficiency of the *RTS1* barrier depends on the expression level of *rtf1*^+^. **A**. The endogenous *rtf1*^+^ promoter results in 70% barrier efficiency. Left: Selected outputs of the model for different efficiencies of delay are compared to the experimental data. Right: Error between the experimental data and the model output (5% steps) measured by the Euclidean distance between the two signals. **B**. adh1*-rtf1* constitutive overexpression results in 90% barrier efficiency. Left: Selected outputs of the model for different efficiencies of delay compared to the experimental data. Right: Error between the experimental data and the model output (5% steps) measured by the Euclidean distance between the two signals.

**Figure S6.**
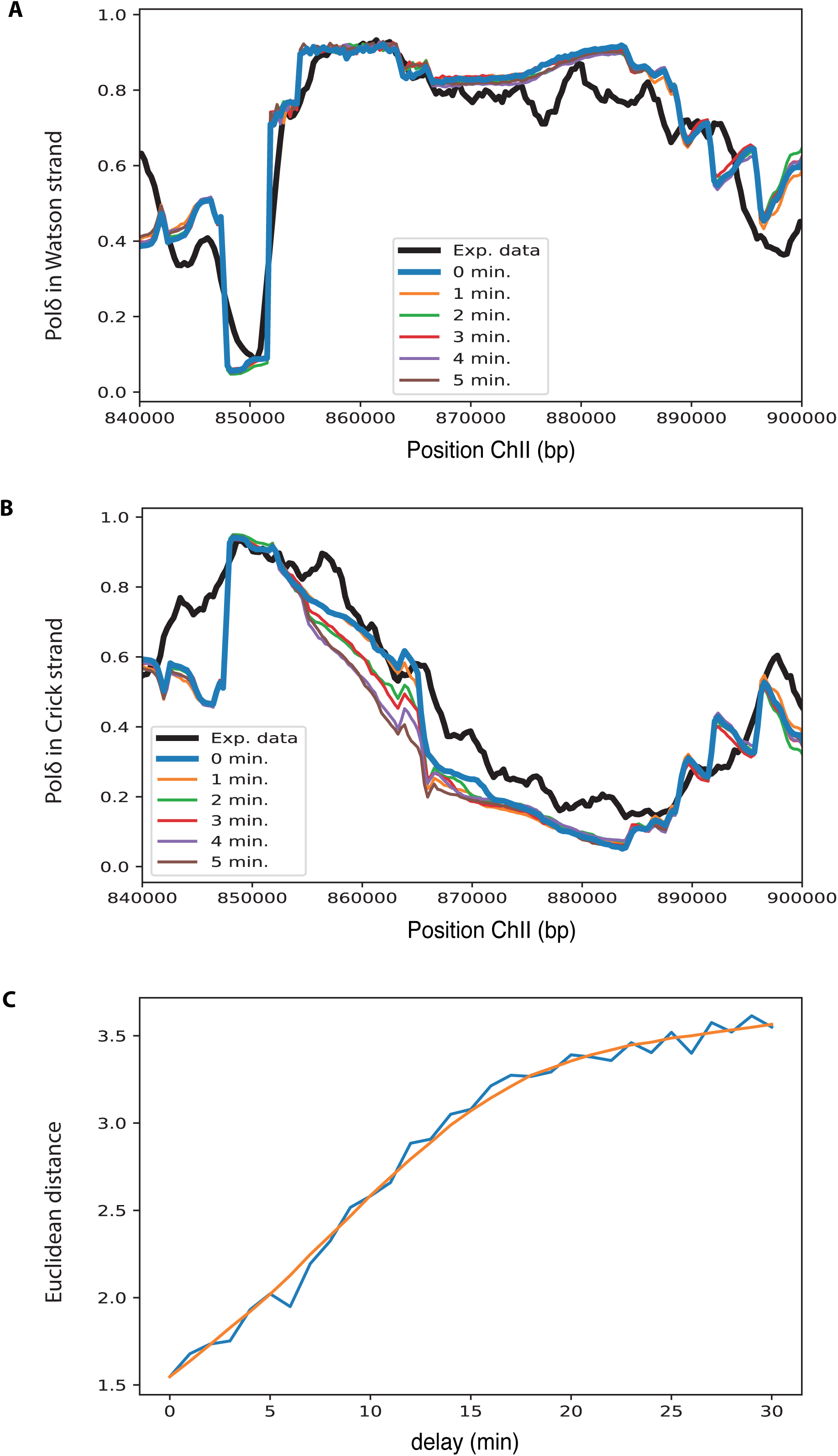
Delay at second *RTS1* barrier after HR restart at the first *RTS1*. Using the replication model, different delays for replication of HR-restarted forks at the second *RTS1* barrier (between 0 and 30 minutes, 1 minute steps) were fitted to the experimental data. **A**. Selected outputs from the model for usage of Pol δ on the Watson strand compared to the experimental data. Note: both when replicated by left-right HR restarted forks, or when replicated by right-left converging canonical forks, the Watson strand is replicated by Pol δ. **B**. Selected outputs from the model for usage of Pol δ on Crick strand compared to the experimental data. Note: A decrease in Pol δ usage on the Crick strand reflects replication by Pol ε from right-left converging forks. Thus, because an increased delay results in more replication by converging canonical forks, this is reflected by a decrease in Pol δ usage. **C**. Error between the experimental data and the model output measured by the Euclidean distance between the two signals; orange curve is a smoothed representation of the errors using a Savitzky-Golay filter.

**Figure S7.**
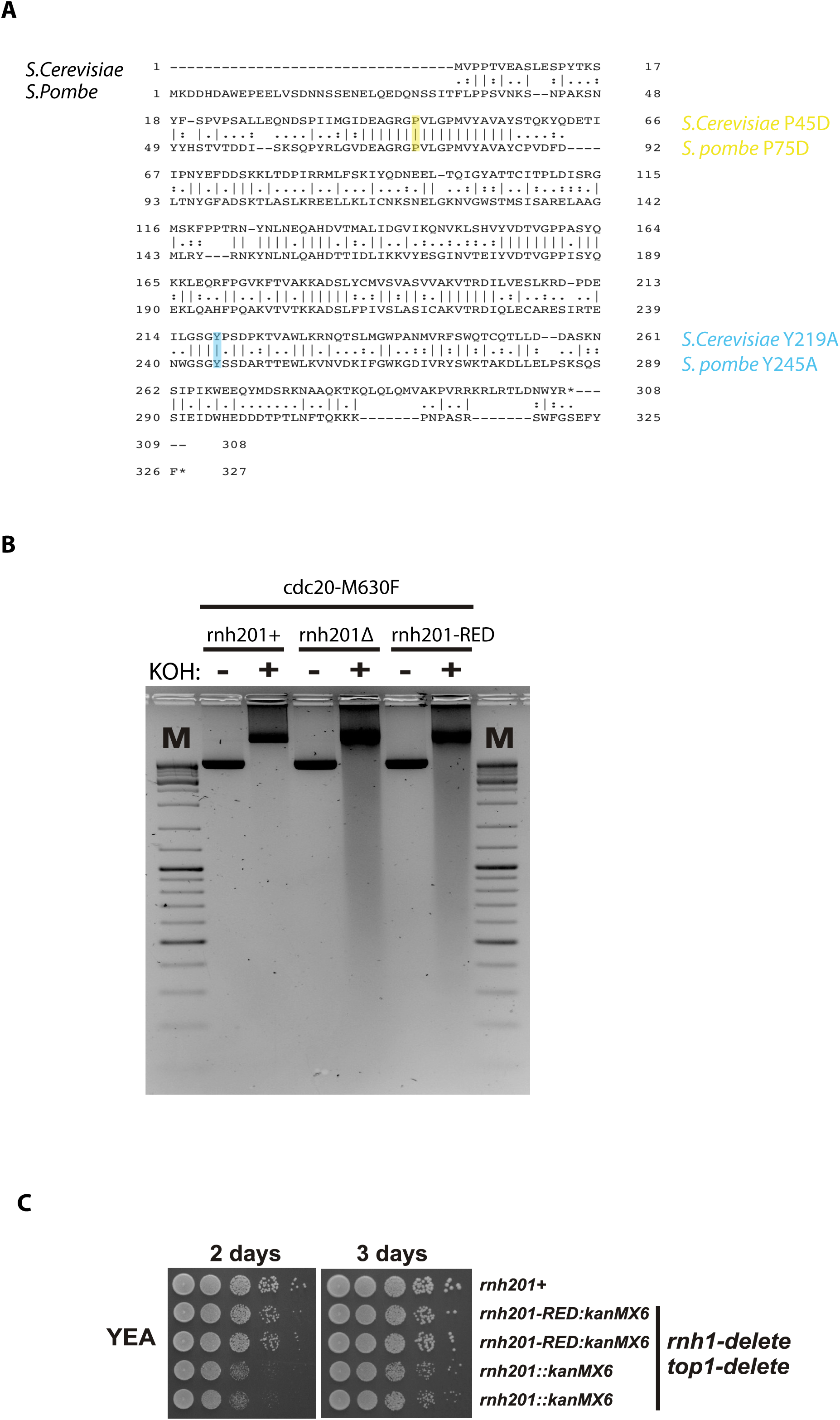
Construction and testing of *rnh201-RED*. In *S. cerevisiae* it has been shown that the synthetic growth defect between the combined *rnh201* and *rnh1* deletions with deletion of *TOP1* is alleviated by restoration of the poly-ribonuclease activity of Rnh201 (i.e. *rnh201-RED*). **A**. Alignment of *S. cerevisiae* and *S. pombe* Rnh201 with highlighted amino acid changes. **B**. Relative DNA fragmentation at incorporated rNTPs after alkaline treatment (for details see the methods section). **C**. Spot test: *top1-d rnh1-d rnh201-d* grow slower than the wild type control (*rnh201*^+^) and *top1-d rnh1-d rnh201-RED*.

**Figure S8.**
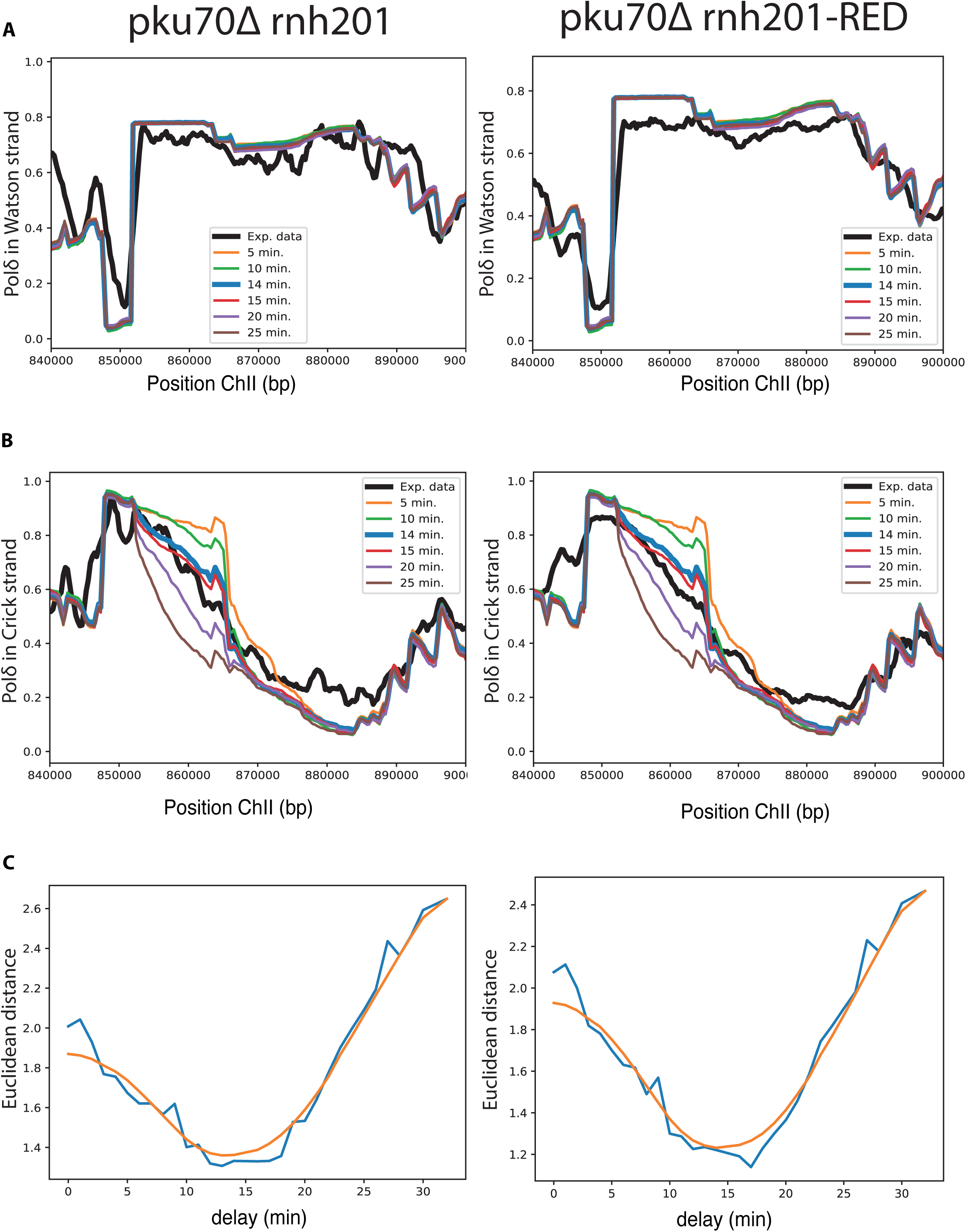
Delay at *RTS1* before HR-restart in absence of Pku70. Using the replication model different delay times of replication at the *RTS1* barrier before HR-dependent restart for the *pku70-d* background were fitted (30 minutes, 1 minute intervals) and compared to the experimental data. The best fit is a 14 minute delay. **A**. Selected outputs from the model for usage of Pol δ on Watson strand compared to the experimental data. Left panel: *pku70-d rnh201-d*. Right panel: *pku70-d rnh201-RED*. Note: both when replicated by left-right HR restarted forks, or when replicated by right-left converging canonical forks, the Watson strand is replicated by Pol δ. **B**. Selected outputs from the model for usage of Pol δ on Crick strand compared to the experimental data. Left panel: *pku70-d rnh201-d*. Right panel: *pku70-d rnh201-RED*. Note: A decrease in Pol δ usage on the Crick strand reflects replication by Pol ε from right-left converging forks. Thus, because an increased delay results in more replication by converging canonical forks, this is reflected by a decrease in Pol δ usage. **C**. Error between the experimental data and the model output measured by the Euclidean distance between the two signals; orange curve is a smoothed representation of the errors using a Savitzky-Golay filter. Left panel: *pku70-d rnh201-d*. Right panel: *pku70-d rnh201-RED*.

**Supplementary Table 1.** *S. pombe* strains.

| Strain | Genotype | Mating type |
| --- | --- | --- |
| 503 | *ade6-704 leu1-32 ura4-d18* | h+ |
| 995 | *cdc6-L591G ade6-704 leu1-32 ura4-d18* | h+ |
| 997 | *cdc20-M630F ade6-704 leu1-32 ura4-d18* | h+ |
| KA219 | *pol1-L850F ade6-704 leu1-32 ura4-d18* | h+ |
| KA217 | *nat:adh1-rtf1 ChII-8535:Rura-10xrRFB RTS1natural::Phleo smt0 cdc2asM17 ade6-704 leu1-32 ura4-D18* | h- |
| KA226 | *ChII-8535:Rura-10xrRFB rtf1::nat RTS1natural::Phleo smt0 cdc2asM17ade6-704 leu1-32 ura4-D18,* | h- |
| KA258 | *rpa3-GFP:kan ChII-8535:Rura-10xrRFB RTS1natural::Phleo rnh201::kan rtf1::nat smt0 rnh1::hyg cdc2asM17 ade6-704 leu1-32 ura4-D18* | h- |
| KA259 | *rpa3-GFP:kan ChII-8535:Rura-10xrRFB RTS1natural::Phleo rnh201::kan nat:adh1-rtf1 rnh1::hyg cdc2asM17 ade6-704 leu1-32 ura4-D18* | h- |
| KA261 | *rpa3-GFP:kan ChII-8535:Rura-10xrRFB RTS1natural::Phleo rtf1::NAT cdc2asM17 ade6-704 leu1-32 ura4-D18* | h- |
| KA262 | *rpa3-GFP:kan ChII-8535:Rura-10xrRFB RTS1natural::Phleo nat:adh1-rtf1 cdc2asM17 ade6-704 leu1-32 ura4-D18* | h- |
| 1144 | *rtf1::nat ChI-3220:kan ade6-704 leu1-32 ura4-d18* | h+ |
| 1145 | *rtf1::nat ChI-3220:kan ade6-704 leu1-32 ura4-d18* | h+ |
| 1146 | *rtf1::nat ChI-4740:kan ade6-704 leu1-32 ura4-d18* | h+ |
| KA2 | *rtf1::nat ChII-8535:kan ade6-704 leu1-32 ura4-d18* | h+ |
| 1158 | *rtf1::nat ChI-3220:TuraR ade6-704 leu1-32 ura4-d18* | h+ |
| 1162 | *rtf1::nat ChI-4740:TuraR ade6-704 leu1-32 ura4-d18* | h+ |
| 1166 | *rtf1::nat ChI-5234:TuraR ade6-704 leu1-32 ura4-d18* | h+ |
| KA7 | *rtf1::nat ChI-8535:TuraR ade6-704 leu1-32 ura4-d18* | h+ |
| 1170 | *rtf1::nat ChI-3220:TuraR rnh201::kan ade6-704 leu1-32 ura4-d18* | h+ |
| 1172 | *rtf1::nat ChI-4740:TuraR rnh201::kan ade6-704 leu1-32 ura4-d18* | h+ |
| 1174 | *rtf1::nat ChI-5234:TuraR rnh201::kan ade6-704 leu1-32 ura4-d18* | h+ |
| KA15 | *rtf1::nat ChI-8535:TuraR rnh201::kan ade6-704 leu1-32 ura4-d18* | h+ |
| 1177 | *ChI-3220:TuraR rnh201::kan cdc6-L591G ade6-704 leu1-32 ura4-d18* | h- |
| 1179 | *ChI-3220:TuraR rnh201::kan cdc20-M630F ade6-704 leu1-32 ura4-d18* | h- |
| 1185 | *ChI-5234:TuraR rnh201::kan cdc6-L591G ade6-704 leu1-32 ura4-d18* | h- |
| 1187 | *ChI-5234:TuraR rnh201::kan cdc20-M630F ade6-704 leu1-32 ura4-d18* | h- |
| KA31 | *ChII-8535:TuraR rnh201::kan cdc6-L591G ade6-704 leu1-32 ura4-d18* | h- |
| KA33 | *ChII-8535:TuraR rnh201::kan cdc20-M630F ade6-704 leu1-32 ura4-d18* | h- |
| KA218 | *rtf1::nat ChII-8535:kan ura4 ade6-704 leu1-32 ura4-d18* | h+ |
| KA233 | *rtf1::nat ChII-8535:kan second RTS barrier ade6-704 leu1-32 ura4-d18* | h+ |
| KA237 | *rtf1::nat ChII-8535:Rura second RTS barrier ade6-704 leu1-32 ura4-d18* | h+ |
| KA238 | *ChII-8535:Rura RTS1natural::Phleo loxP-rnh201-RED-kanR-loxM3 nat:adh1-rtf1 ade6-704 leu1-32 ura4-D18* | h- |
| KA156 | *ChII-8535:Rura-10xrRFB RTS1natural::Phleo rnh201:kan nat:adh1-rtf1 ade6-704, leu1-32 ura4-D18* | h- |
| KA199 | *ChII-8535:Rura-10xrRFB cdc6-L591G RTS1natural::Phleo rnh201:kan nat:adh1-rtf1 ade6-704 leu1-32 ura4-D18* | h- |
| KA200 | *ChII-8535:Rura-10xrRFB cdc20-M630F RTS1natural::Phleo rnh201:kan nat:adh1-rtf1 ade6-704 leu1-32 ura4-D18* | h- |
| KA241 | *ChII-8535:Rura second RTS barrier RTS1natural:Phleo rnh201-RED:kan nat:adh1-rtf1 ade6-704 leu1-32 ura4-D18* | h- |
| KA256 | *cdc20-M630F ChII-8535:Rura second RTS barrier RTS1natural::Phleo rnh201-RED:kan nat:adh1-rtf1 ade6-704 leu1-32 ura4-D18* | h- |
| KA260 | *cdc6-L591M ChII-8535:Rura second RTS barrier RTS1natural::Phleo rnh201-RED:kan nat:adh1-rtf1 ade6-704 leu1-32 ura4-D18* | h- |
| 1048 | *rnh201::kan ade6-704 leu1-32 ura4-d18* | h+ |
| KA103 | *ChII-8535:tandemRTS1-ura4-10xrRFB rtf1::nat rnh201::kan ade6-704 leu1-32 ura4-D18* | h+ |
| SAL445 | *RTS1natural::Phleo rtf1:nat smt0* | h- |
| KA107 | *ChII-8535:tandemRTS1-ura4-10xrRFB rtf1::nat rnh201::kan RTS1natural::Phleo smt0 ade6-704 leu1-32 ura4-D18* | h- |
| KA169 | *cdc6-L591G ChII-8535:tandemRTS1-ura4-10xrRFB rnh201:kan RTS1natural::Phleo smt0 ade6-704 leu1-32 ura4-D18* | h- |
| KA170 | *cdc20-M630F ChII-8535:tandemRTS1-ura4-10xrRFB rnh201::kan RTS1natural::Phleo smt0 ade6-704 leu1-32 ura4-D18* | h- |
| KA76 | *Ku70:leu2 ChII-8535:Rura-10xrRFB RTS1natural::Phleo smt0x rnh201::kan rtf1::nat ade6-704 leu1-32 ura4d18* | h- |
| KA77 | *Ku70:leu2 ChII-8535:Rura-10xrRFB RTS1natural::Phleo smt0x rnh201::kan ade6-704 leu1-32 ura4d18* | h- |
| KA78 | *cdc6-L591G Ku70:leu2 ChII-8535:Rura-10xrRFB RTS1natural::Phleo smt0x rnh201::kan rtf1::nat ade6-704 leu1-32 ura4d18* | h- |
| KA79 | *cdc20-M630F Ku70:leu2 ChII-8535:Rura-10xrRFB RTS1natural::Phleo smt0x rnh201::kan rtf1::nat ade6-704 leu1-32 ura4d18* | h- |
| KA80 | *cdc6-L591G Ku70:leu2 ChII-8535:Rura-10xrRFB RTS1natural::Phleo smt0x rnh201::kan ade6-704 leu1-32 ura4d18* | h- |
| KA81 | *cdc20-M630F Ku70:leu2 ChII-8535:Rura-10xrRFB RTS1natural::Phleo smt0x rnh201::kan ade6-704 leu1-32 ura4d18* | h- |
| KA156 | *ChII-8535:Rura-10xrRFB RTS1natural::Phleo rnh201::kan nat:adh1-rtf1 ade6-704 leu1-32 ura4-D18* | h- |
| BAY125 | *cdc6-L591G ChII-8535:Rura-10xrRFB RTS1natural::Phleo rnh201::kan rtf1::nat ade6-704 leu1-32 ura4-D18* | h- |
| BAY123 | *cdc20-M630F ChII-8535:Rura-10xrRFB RTS1natural::Phleo rnh201::kan rtf1::nat ade6-704 leu1-32 ura4-D18* | h- |
| BAY126 | *cdc6-L591G ChII-8535:Rura-10xrRFB RTS1natural::Phleo, rnh201::kan, ade6-704 leu1-32 ura4-D18* | h- |
| BAY124 | *cdc20-M630F ChII-8535:Rura-10xrRFB RTS1natural::Phleo rnh201::kan ade6-704 leu1-32 ura4-D18* | h- |
| KA161 | *ChII-8535:Rura-10xrRFB rnh201-RED:kan ade6-704 leu1-32 ura4-D18* | h- |
| KA164 | *ChII-8535:Rura-10xrRFB rnh201-RED:kan RTS1natural::Phleo rtf1::nat ade6-704 leu1-32 ura4-D18* | h- |
| KA171 | *cdc6-L591G ChII-8535:Rura-10xrRFB rnh201-RED:kan RTS1natural::Phleo, smt0 ade6-704 leu1-32 ura4-D18* | h- |
| KA172 | *cdc6-L591G ChII-8535:Rura-10xrRFB rnh201-RED:kan RTS1natural::Phleo, rtf1::nat smt0 ade6-704 leu1-32 ura4-D18* | h- |
| KA173 | *cdc20-M630F ChII-8535:Rura-10xrRFB rnh201-RED:kan RTS1natural::Phleo smt0 ade6-704 leu1-32 ura4-D18* | h- |
| KA174 | *cdc20-M630F ChII-8535:Rura-10xrRFB rnh201-RED:kan RTS1natural::Phleo, rtf1::nat smt0 ade6-704 leu1-32 ura4-D18* | h- |
| KA203 | *ChII-8535:Rura-10xrRFB pol1-L850F rnh201::kan RTS1natural::Phleo ade6-704 leu1-32 ura4-D18* | h- |
| KA204 | *ChII-8535:Rura-10xrRFB pol1-L850F rnh201-RED:kan RTS1natural::Phleo, ade6-704 leu1-32 ura4-D18* | h- |
| KA205 | *ChII-8535:Rura-10xrRFB pol1-L850F rnh201::kan rtf1::nat RTS1natural::Phleo ade6-704 leu1-32 ura4-D18* | h- |
| KA206 | *ChII-8535:Rura-10xrRFB pol1-L850F rnh201-RED:kan rtf1::nat RTS1natural::Phleo ade6-704 leu1-32 ura4-D18* | h- |
| KA215 | *ChII-8535:Rura-10xrRFB rnh201-RED:kan Ku70::hyg RTS1natural::Phleo smt0 rtf1::nat ade6-704 leu1-32 ura4-D18* | h- |
| KA220 | *cdc6-L591G ChII-8535:Rura-10xrRFB rnh201-RED:kan Ku70::hyg RTS1natural::Phleo smt0 rtf1::nat ade6-704 leu1-32 ura4-D18* | h- |
| KA221 | *cdc6-L591G ChII-8535:Rura-10xrRFB rnh201-RED:kan Ku70::hyg RTS1natural::Phleo smt0 ade6-704 leu1-32 ura4-D18* | h- |
| KA222 | *cdc20-M630F ChII-8535:Rura-10xrRFB rnh201-RED:kan Ku70::hyg RTS1natural::Phleo smt0 rtf1::nat ade6-704 leu1-32 ura4-D18* | h- |
| KA223 | *cdc20-M630F ChII-8535:Rura-10xrRFB rnh201-RED:kan Ku70::hyg RTS1natural::Phleo smt0 ade6-704 leu1-32 ura4-D18* | h- |
| BAY4 | *ChI-3220:10xrRFB ade6-704 leu1-32 ura4-D18* |  |
| BAY12 | *cdc6-L591G ChI-3220:10xrRFB rnh201:kan ade6-704 leu1-32 ura4-D18* | h- |
| BAY16 | *cdc20-M630F ChI-3220:10xrRFB rnh201:kan rtf1::nat ade6-704 leu1-32 ura4-D18* | h- |
| KA39 | *rtf1::nat ChII-8535:kan-ura4 ade6-704 leu1-32 ura4-d18* | h+ |
| KA56 | *rtf1::nat ChII-8535:kan-10xrRFB ade6-704 leu1-32 ura4-d18* | h+ |
| KA58 | *rtf1::nat ChII-8535:Rura-10xrRFB ade6-704 leu1-32 ura4-d18* | h+ |
| AW1480 | *rnh201::ura4 ura4D18 leu1-32* | h- |
| AW1563 | *rnh201-RED:kan ura4D18 leu1-32* | h- |
| AW1565 | *rnh201-RED:hyg ura4D18 leu1-32* | h- |
| AW1566 | *rnh201-RED:nat ura4D18 leu1-32* | h- |
| AW1466 | *cdc20M630F ade6-704 ura4D18 leu1-32* | h- |
| AW1405 | *cdc20M603F rnh201::kan ade6-704 leu1-32 ura4D18* | h- |
| AW1232 | *rnh201-RED:kan cdc20-M630F* | h- |
| AW278 |  | h- |
| AW1229 | *rnh201-RED:kan rnh1::nat top1::hyg* | h- |
| AW1230 | *rnh201-RED:kan rnh1::nat top1::hyg* | h+ |
| AW1198 | *rnh201::kan rnh1::nat top1::hyg* | h+ |
| AW1199 | *rnh201::kan rnh1::nat top1::hyg* | h+ |

**Supplementary Table 2.**
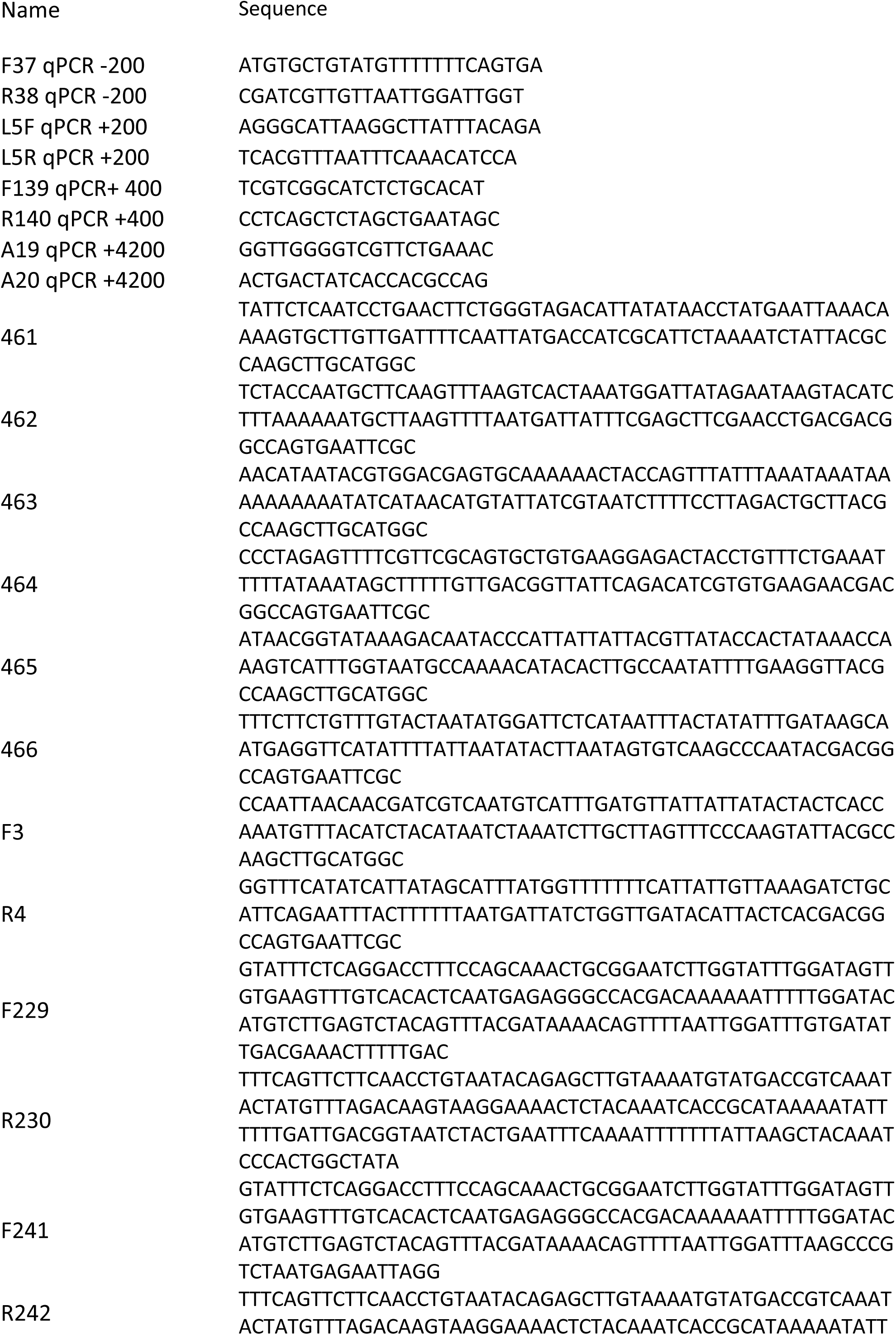
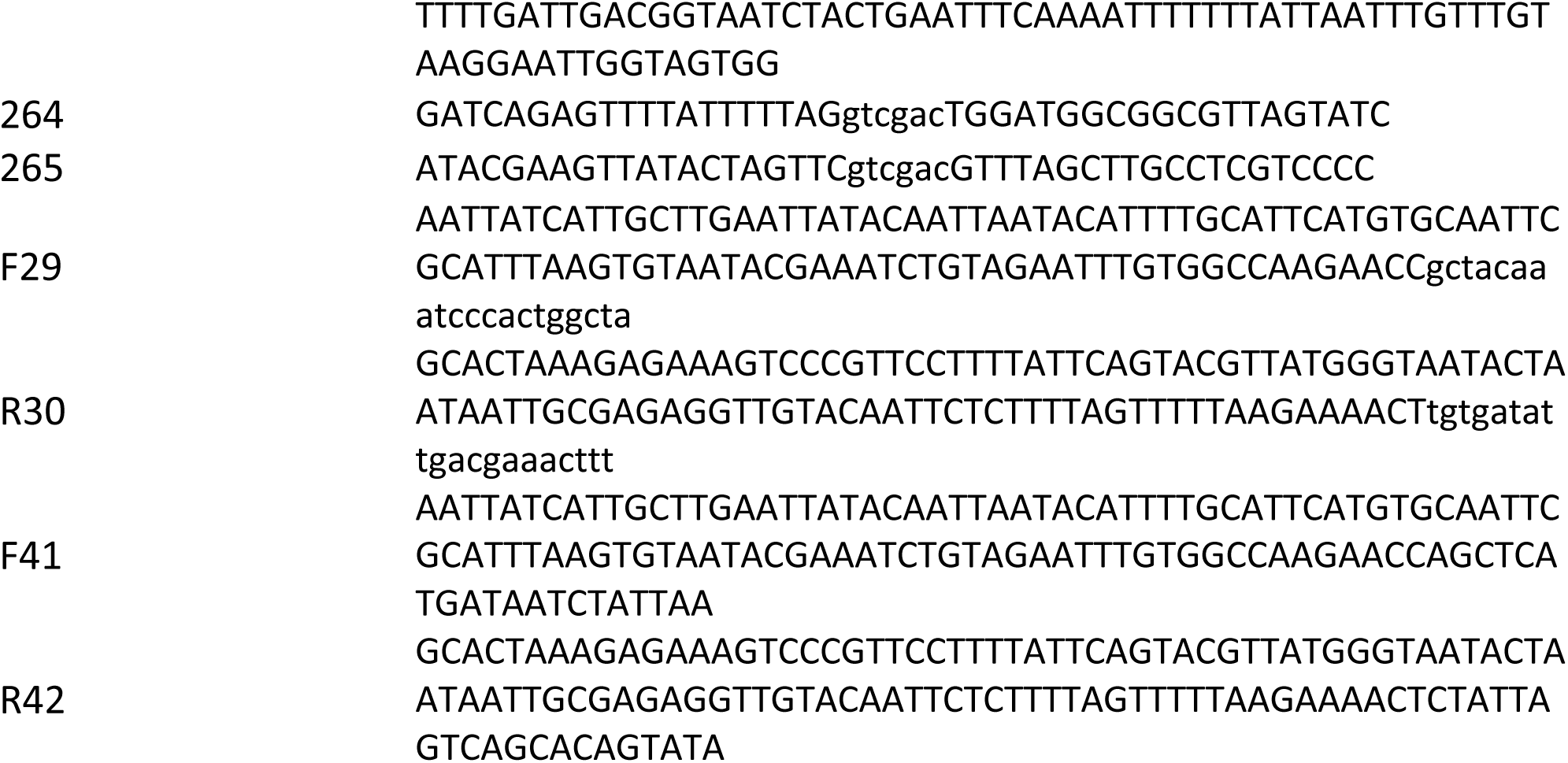
Primers.

## Online Materials and Methods

### Strain construction

To introduce the *RTS1* sequence into 4 loci analysed in Figure S1, the kan cassette from pFAX-kanMX6 was amplified by the following primers: 461/462 (*mns1* locus), 463/464 (*tel2* locus), 465/466 (*mek1* locus), F3/R4 (*spo9* locus). PCR amplified fragments (which introduce a loxP and a loxM site flanking the kan cassette) were transformed into strain 503 to form strains 1144, 1146, 1148 and KA2 respectively. These strains were subsequently transformed with pAW8-RTS1-ura-2/3ter^10^ to integrate a construct between the lox sites containing RTS1-ura-2/3ter. This resulted in strains 1158, 1162, 1166, KA7

To create the BAY4 strain used to analyse the effect of the rRFB (Figure S2) strain 1144 (containing loxP-loxM flanking the kan cassette on ChrI) was transformed with pAW8-10xrRFB-ura to integrate 10xRFB-ura between the lox sites.

To create the optimised ChrII locus (Figure 1) the 10xrRFB sequence was integrated downstream of *RTS1* barrier. Primers F29 and R30 were used to amplify *ura4*^+^ and the product was transformed into KA2 and checked for insertion onto the ChrII between *SPBC36*.*10*.*1* and *SPBC36*.*11*.*1* to form strain KA39. The *ura4*^+^ gene was then replaced by *10xrRFB* using a fragment amplified from pAW8-10xrRFB using primers F41 and R42 to form strain KA56. KA56 was transformed by pAW8-RTS-ura including loxP and loxM sites. This created the strain KA58.

To construct the inverted *RTS1* locus used in Figure 2, primers F229 and R230 were used to amplify *ura4*^+^ and the product was transformed into KA2 and checked for insertion onto the ChrII between *pwp2* and *wdr83* to form strain KA218. The *ura4*^+^ gene was then replaced by *RTS1* using a fragment amplified with primers F241 and R242 to form strain KA233. KA233 was transformed by pAW8-RTS-ura including loxP and loxM sites. The kan cassette between *spo9* and *clr6* on ChrII close to early firing origin was replaced by the *RTS1* construct. This created the strain KA237.

To construct the tandem *RTS1* barrier used in Figure 3 we modified pAW8-RTS-ura to inserted a second *RTS1* downstream of *ura4*^+^ between the *Sac*I-*Sal*I sites. The plasmid was transformed into KA56 which contains the kan cassette flanked by loxP and LoxM sites. Cre-lox recombination replaced the kan cassette by the RTS-ura-RTS cassette between *spo9* and *clr6* on ChrII, to give strain KA101.

To construct *rnh201-RED* mutant allele the *S. cerevisiae* Rnh201-RED mutant sequence^26^ was aligned to *S. pombe* Rnh201. *S. cerevisiae* Rnh201 P45D and Y219A changes corresponded to *S. pombe* Rnh201 P75D and Y245A (FigS1A). The *S. pombe rnh201-P75D-Y245A* gene sequence was synthesised (Integrated DNA technologies) and cloned into Cre-expression vector pAW8^31^ to create rnh201-RED-pAW8. The hphMX6, natMX6 and kanMX6 antibiotic resistance markers^32^ were amplified using primers 264 and 265 and cloned into rnh201-RED-pAW8 to create rnh201-RED-hphMX6-pAW8, rnh201-RED-natMX6-pAW8 and rnh201-RED-kanMX6-pAW8 respectively. All constructs were confirmed by sequencing. The rnh201-RED-hphMX6-pAW8, rnh201-RED-natMX6-pAW8 and rnh201-RED-kanMX6-pAW8 plasmids were used to transform *S. pombe*. ‘rnh201 base strain’ AW1480 and the rnh201-RED-MX6 sequences integrated by recombinase-mediated cassette exchange^31^. This created strains AW1563 (*rnh201-RED:hphMX6*), AW1565 (r*nh201-RED:natMX6*) and AW1566 (*rnh201-RED:kanMX6*).

To test rNTP incorporation rates, strain AW1563 (*rnh201-RED:hphMX6*) was crossed to strain AW1227 (*cdc20M630F*) to create strain AW1232 (*rnh201-RED:kanMX6, cdc20M630F*). Control strains included *rnh201*^+^, AW1466 (*cdc20-M630F*) and AW1405 (*rnh201Δ, cdc20-M630F*). Approximately 3×10^8^ logarithmically growing cells were collected and the genomic DNA extracted^33^ and incubated in the presence of 0.3N KOH for 1hour at 55°C. The alkali was then neutralised by the addition of equimolar amount of HCl and the reaction mix isopropanol precipitated. A portion of the genomic DNA was untreated to confirm the integrity of the DNA extract. Following 2% agarose/1xTBE gel electrophoresis and staining with acrydine orange (Invitrogen), the degree of fragmentation observed was compared relative to the non-treated DNA control.

To perform spot assays of *S. pombe* strains AW278 (WT), AW1229/AW1230 (*rnh201-RED, rnh1-d top1-d*) and AW1198/AW1199 (*rnh201-d, rnh1-d* and *top1-d*) were serially diluted 10-fold in water and spotted on YEA plates.

A list of strains and associated genotypes is given in Table S1.

### Cell synchronisation

Cells were synchronised using the *cdc2-asM17* allele^34^ and 3-Br-PP1 (2uM final, Abcam) at 28°C for 3 hours. Upon the removal of 3-Br-PP1, cells were grown at 30°C and samples collected at the indicated times.

### ChIP-qPCR

40 ml of log-phase cells were grown in YES or EMM liquid media at 30°C and cross-linked in 1% formaldehyde (15min RT°) followed by addition of 5ml 2.5 M glycine for 5 min. Cells were centrifuged (4000g, 5min) and washed with 10ml PBS and the pellet frozen. For RNA:DNA hybrid ChIP cells were not crosslinked but NaN3 (0.1% final) was added.

Cell pellets were resuspended in 400ul ChIP lysis buffer (50 mM HEPES pH 7.4; 140 mM NaCl; 1 % Triton X100; 0.1 % NaDeoxycholate; protease inhibitors), disrupted by glass bead beating using a FastPrep-24 (MPbiomedical) homogenizer. Homogonised material was harvested by 1-minute centrifugation to remove glass beads, washed once with 1 ml ChIP lysis buffer and resuspended in 350 ml of ChIP lysis buffer. The chromatin DNA was sheared by a Qsonica ultrasonicator to the size range of approximately 500bp. Samples were centrifuged for 1 minute and the supernatant incubated with antibody (GFP polyclonal antibody from Invitrogen, cat.no. A-11122) or S9.6 (Sigma-Aldrich, cat.no. MABE1095) for 1 hour at 4°C. Protein G coupled Dynabeads (Thermo Fisher) were used for immunoprecipitation overnight at 4°C. The beads were washed twice for 5 min at 4°C in 1 ml of each of the following buffers: ChIP lysis buffer, high salt lysis buffer (50 mM HEPES pH 7.4; 500 mM NaCl; 1 % Triton X100; 0.1 % NaDeoxycholate) and wash buffer (10 mM Tris pH 8.0; 250 mM LiCl; 0.5 % NP-40; 0.5 % NaDeoxycholate; 1 mM EDTA) and once in TE buffer (10 mM Tris pH 8.0, 1 mM EDTA) for a minute. Beads were resuspended in (110 ul) elution buffer (50 mM Tris pH 8.0; 1 % SDS; 10 mM EDTA) and 3ul cocktail of Rnases (Invitrogen) was added and incubated at 30°C for 2 hours with shaking. 5 ul of 20mg/ml Proteinase K (Sigma Aldrich) was added and incubated at 65°C for 2 hours with shaking. The samples were purified using a PCR purification kit (Quiagen) and analysed by qPCR using LUNA qPCR mix (NEB) on a real-time detection system (Agilent). Fold enrichment was calculated as ChIP/Input. Primers used for qPCR are listed in Table S2.

### Polymerase usage sequencing (Pu-seq)

The published protocol^19^ was used with minor modifications: size selection was performed using a Blue Pippin (Sage Science). In some instances we used *rnh201-RED* instead of *rnh201::kan* (see results).

### Mathematical modelling

We simulated the replication process using a variation of the model presented by Kelly and Callegari^18^. We extended the model to include time- and location-dependent fork velocity to allow the modelling of the replication barriers. The model determines origin locations and efficiency based on the AT-richness and transcription activity. We optimised the parameters of the model to globally fit the wild type replication profile, using the line search approach akin to the methodology of Kelly and Callegari. We suppressed some minor origins (which figure 2 demonstrates are not used) in the region of interest to fit to the traces of polymerase usage in the wild type strain. These origins were otherwise present in the simulation but not in acquired data by Pu-seq.

We modelled the fork barrier by means of the fork velocity: at the location of the barrier, the modelled fork velocity drops to zero for some time, and then increases to the original value. We use a constant velocity of 1800 kb/min for the rest of the chromosome. We defined separately the velocity of left- and right-travelling forks in order to model unidirectional fork barriers.

We used the Euclidean norm of the difference between model solution and experimental data to quantify the model error. Where appropriate a Savitzky-Golay filter was used to smooth the line. In all the simulations we used an ensemble of 1000 cells, and then we validated the optimal result by further simulating an ensemble of 10000 cells.

All the quantities that we optimised using the model were found by performing a line search. We used this approach to find the model predictions for the fraction of forks arrested at the *RTS1* barrier, the time to restart for arrested forks and the effect of the *RTS1* barrier on restarted forks.

